# *Streptococcus mitis* bacteriocins drive contact-dependent lysis of *S. pneumoniae* facilitating transformation in multispecies environments

**DOI:** 10.1101/2025.08.25.672098

**Authors:** João Borralho, João Lança, Joana Bryton, Wilson Antunes, Raquel Sá-Leão

## Abstract

Natural competence allows bacterial species like *Streptococcus pneumoniae* and *S. mitis* to acquire environmental DNA, driving horizontal gene transfer (HGT) and adaptation. In *S. pneumoniae*, a human pathogen, competence-induced predation is well characterized and involves the release of bacteriocins and a murein hydrolase to lyse noncompetent siblings and liberate DNA. In contrast, in the human commensal *S. mitis*, mechanisms mediating DNA acquisition remain poorly understood. Here, we identify a diverse set of competence-associated bacteriocins (*cab*) that are produced by *S. mitis* during the late phase of competence. We focus on one bacteriocin pair, CabAB, that triggers contact-dependent growth inhibition and lysis of *S. pneumoniae* through activation of the major pneumococcal autolysin LytA. We demonstrate that CabAB compromises *S. pneumoniae* membrane integrity, leading to formation of intracellular membrane aggregates and the release of cytoplasmatic content, thereby increasing available DNA, which enhances HGT from *S. pneumoniae* to *S. mitis* in biofilms. These findings uncover a mechanism of interspecies predation and gene acquisition, revealing a critical role for competence-associated bacteriocins in shaping evolutionary dynamics of streptococci.

**Importance:** Many streptococci are naturally competent, acquiring environmental DNA through transformation. This includes pathogens like *S. pneumoniae* and commensals like *S. mitis,* which can exchange genetic material through horizontal gene transfer (HGT). For example, *S. mitis* can acquire pneumococcal capsules, leading to its misidentification in polymicrobial samples such as those obtained from the upper respiratory tract. Understanding the drivers of HGT between these species is therefore critical. Here, we characterize a competence-induced bacteriocin cluster in *S. mitis*. These bacteriocins lyse pneumococci, promoting DNA release and enhancing gene transfer in dual-species biofilms. Our findings uncover a mechanism by which competence-associated predation promotes interspecies HGT, shaping the evolution and epidemiology of streptococcal populations.

## Introduction

Natural competence for genetic transformation is a developmental program found in certain bacterial species that enables the uptake of naked DNA from the environment. Once internalized, this DNA can be incorporated into the genome through homologous recombination, making natural transformation one of the main modes of horizontal gene transfer (HGT). This phenomenon was first described in *Streptococcus pneumoniae* (or pneumococcus), a species capable of exchanging pathogenic traits such as the capsule by taking up DNA from other pneumococci (1).

In *S. pneumoniae*, competence is a tightly regulated transient physiological state which is governed by a cell-cell signaling system. The competence-stimulating peptide (CSP), encoded by *comC*, is processed and exported via the ATP-binding cassette transporter ComAB. Extracellular CSP is sensed by the membrane-bound histidine kinase ComD (2–4), which undergoes autophosphorylation triggering subsequent phosphorylation of the response regulator ComE (5). Phosphorylated ComE increases expression of the early competence cascade including *comAB* and *comCDE*, creating a positive feedback loop. It also induces expression of the two copies of the alternative sigma factor (*sigX1* and *sigX2*), marking the onset of the late phase of competence (6).

In the late phase of competence, expression of all genes required for DNA uptake, processing and recombination – the transformasome – is induced (7). Here, pneumococci produce the competence-induced bacteriocins A and B (CibAB), along with the immunity protein CibC, which provides self-immunity to CibAB (8). Because competence may not be synchronized across the whole population (9–11), CibAB release on solid media may lead to lysis of genetically identical (kin) noncompetent cells that have not yet expressed *cibC*. Additionally, expression of the murein hydrolase CbpD, against which competent cells are protected via ComM expression, is essential for kin predation in liquid cultures. Notably, CbpD has also been implicated in lysis of non-kin, closely related streptococci, thereby expanding the gene pool from which *S. pneumoniae* may acquire DNA (12). CibAB and CbpD have also been associated to competitive interactions *in vivo*, where their expression by colonizing pneumococci provides a competitive advantage over other *S. pneumoniae* invaders (13).

A close relative of the pneumococcus, *S. mitis*, is also naturally transformable. *S. mitis* is a common commensal of the human nasopharynx and oral cavity, where it typically establishes long-term colonization without causing disease (14). Despite close phylogenetic relationship with *S. pneumoniae*, and a strikingly similar competence regulatory network, *S. mitis* exhibits key differences in terms of bacteriocin content. While *S. mitis* typically encodes *cbpD*, most strains lack *cibAB*, suggesting that *S. mitis* may rely on an alternative mechanism for predation during competence (15–17).

Although the frequency at which *S. mitis* acquires DNA from *S. pneumoniae* is not well established, such events have been documented. These include, for example, the acquisition of pneumococcal *pbp2x* mosaic blocks (18–20). Some *S. mitis* have also been found to express some pneumococcal or pneumococcal-like capsules, which can cross-react with antisera against serotypes 1, 5, 17F, 18A, 19C, 21, 39 and 40 (21–24).

A competence-associated bacteriocin (*cab*) cluster, which is absent in *S. pneumoniae*, has been identified in *S. mitis*. This cluster is located upstream of the competence transporter *comAB* and can encode multiple *cab* bacteriocins distinct from CibAB (15, 16, 25–27). However, the role of *cab* in *S. mitis* predation of kin and non-kin, and its contribution to DNA acquisition, remains unexplored.

Here, we characterize the arsenal of *cab* bacteriocins carried by *S. mitis* and provide evidence that some specifically mediate lysis of *S. pneumoniae*. We also show that pneumococcal DNA can be efficiently taken up and recombined by *S. mitis* and that Cab-mediated lysis accelerates gene transfer from *S. pneumoniae* to *S. mitis*. Together, these findings shed light on previous observations of pneumococcal DNA integration in *S. mitis* genomes.

## Results

### *S. mitis* encodes a diverse arsenal of competence-associated bacteriocins (*cab*)

We previously identified and characterized a limited number of *S. mitis* strains encoding multiple competence-associated bacteriocins (*cab*), most of which lacked the pneumococcal bacteriocins *cibAB* and associated immunity protein *cibC* (27). To investigate the broader distribution of *cab* and *cibAB* in *S. mitis*, we surveyed publicly available genomes, leveraging the conserved genomic context of *cab* clusters – located upstream of *comAB* and downstream of *plsX*-*acpP*. This region was identified and extracted from 253 of 332 *S. mitis* genomes (Fig. 1A and Table S1). The remaining genomes had contig breaks and were excluded from further analysis.

**Fig. 1.**
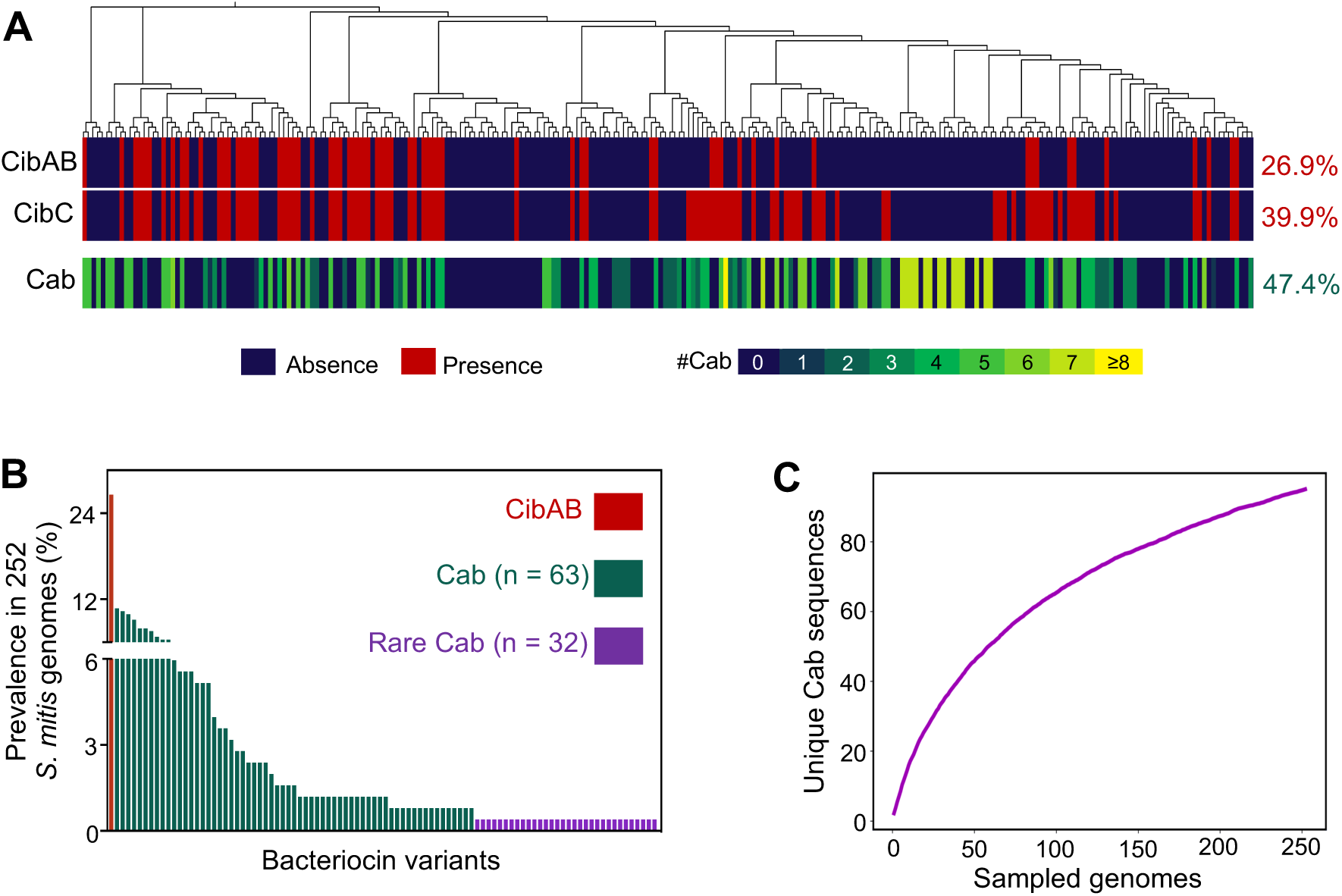
*Streptococcus mitis* genomes encode a plethora of competence-associated bacteriocins (*cab*). (A) Core-SNP phylogeny of 253 *S. mitis* genomes and distribution of competence-induced bacteriocins A and B (*cibAB*), dedicated immunity (*cibC*), and competence-associated bacteriocin (*cab*) genes. (B) Prevalence of CibAB and 95 Cab variants in the *S. mitis* population. (C) Rarefaction analysis of mature Cab amino acid sequences found in the *S. mitis* population.

Among the 253 genomes analyzed, 26.9% harbored *cibABC*, a striking contrast with *S. pneumoniae*, where these genes are universally present. An additional 13.0% of strains carried *cibC* but not *cibAB* – often referred to as “cheater” phenotype (28), in which strains are immune to CibAB-mediated killing but do not produce the bacteriocins themselves. In contrast, 47.4% of *S. mitis* encoded between one to eleven unique *cab* genes that contain a double glycine leader sequence ((M/L/V)XXXXGG), which is recognized by ComAB (29). We defined *cab* bacteriocins as all non-*cib* bacteriocin genes that were found in this particular location. In total, 95 distinct Cab variants were found, which grouped into 28 clusters based on sequence similarity (Fig. 1B and Fig. S1). Overall, 61.3% of *S. mitis* genomes carried at least one *cab* gene or the *cibABC* operon (Table S1).

Cab variants were present at low frequencies (<10%) in the *S. mitis* population, with 32 of 95 bacteriocin variants only being found in a single strain (Fig. 1B). Despite this extensive diversity, rarefaction analysis suggested that Cab diversity is not yet saturated (Fig. 1C); sequencing more *S. mitis* will likely reveal new Cab sequences.

These findings highlight *S. mitis* as a reservoir of highly diverse and largely unexplored competence-associated bacteriocins.

### Cab bacteriocins are expressed in the late phase of competence

*S. mitis* G22, previously described by us, encodes a *cab* operon consisting of two putative bacteriocins and an associated immunity protein (hereafter referred as *cabAB* and *cabC*, respectively; *cabA* belongs to cluster 7 and *cabB* belongs to cluster 21) (Fig. 2A and Fig. S1) (27). This strain was selected to study *cab* regulation in *S. mitis* and its role in competitive behaviors. Both bacteriocins and the immunity protein it encodes are rare (30) and thus unlikely to encounter pre-existing immunity in other strains. Notably, its *cab* promoter contains five putative binding sites for the alternative sigma factors (SigX boxes) (27), suggesting that *cabABC* may be expressed during the late phase of competence.

**Fig. 2.**
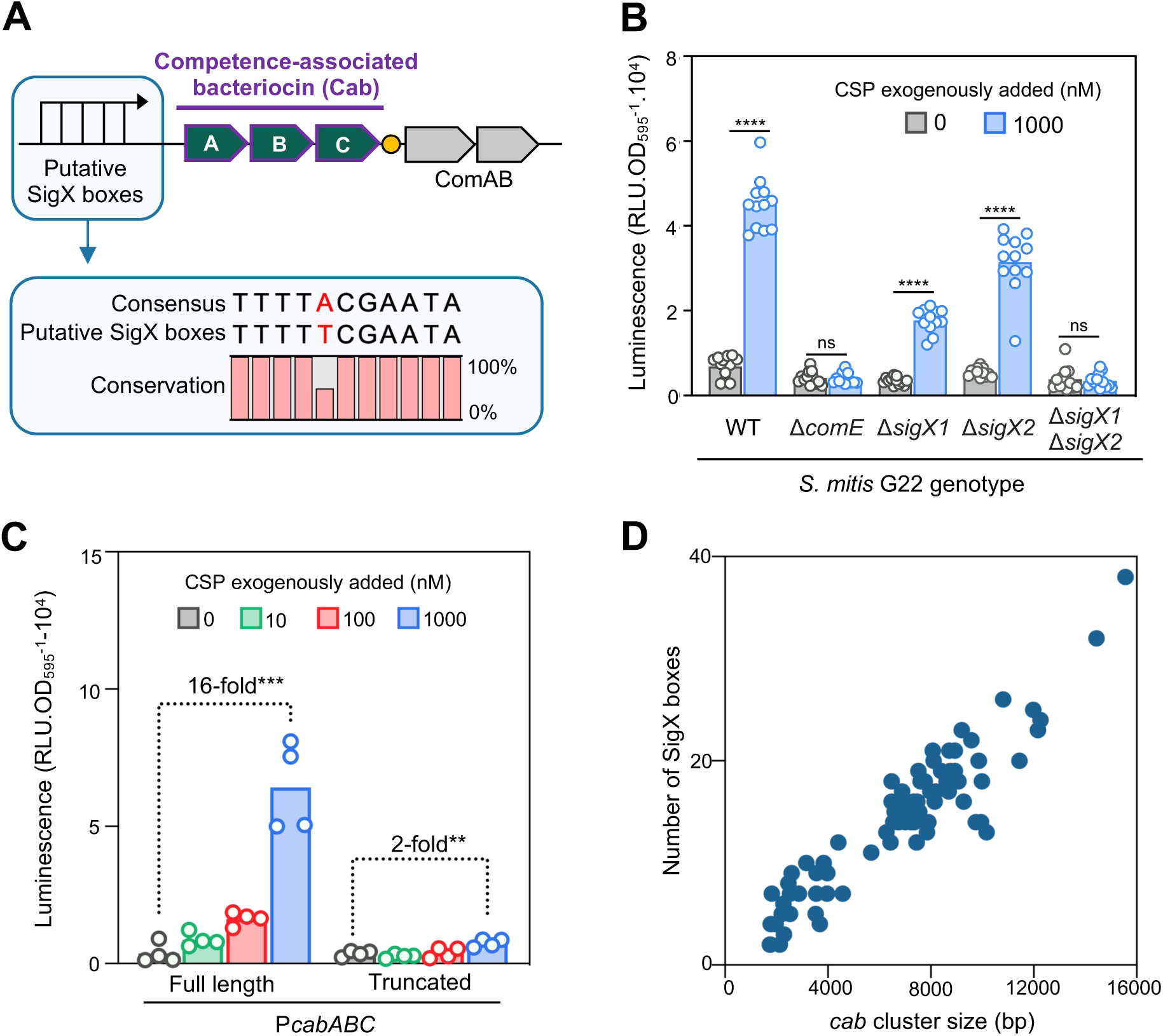
Cab bacteriocins are expressed at the late phase of competence. (A) Genetic organization of *cab* cluster of strain *S. mitis* G22, containing two putative bacteriocin (*cabAB*) and an immunity protein (*cabC*) upstream of the ABC-transporter *comAB* genes. The *cabABC* operon promoter (P*cabABC*) contains five tandem copies of SigX-binding sites (SigX box; SigX is the alternative sigma factor that controls gene expression during the late phase of competence). Expression of *comAB* is driven from a different promoter and induced by ComE (represented as an orange circle). (B) Expression of *cabABC* in *S. mitis* G22 containing a P*cabABC*-luciferase (*luc*) reporter, ectopically inserted in its genome. Expression of *cabABC* was inferred by expression of *luc*. *S. mitis* was pre-grown in acidic medium to avoid natural competence development and synthetic cognate competence-stimulating peptide (CSP_G22_) was added to wild-type *S. mitis* and mutants lacking competence regulators. (C) Dose-response of *cabABC* expression across CSP_G22_ concentrations for the *S. mitis luc* reporter containing either the full-length P*cabABC* promoter (which contains five SigX boxes), or a truncated version containing only the most proximal SigX box. (D) Number of SigX boxes within *cab* clusters of varied sizes. Data in panels B and C represent three to four independent experiments. Unpaired *t*-tests were performed to compare *cab* expression. ** *P* < 0.05, *** *P* < 0.001, **** *P* < 0.0001.

To investigate the dynamics of *cab* expression, the native *cabABC* promoter was ectopically inserted into the *S. mitis* G22 genome, upstream of a firefly luciferase reporter (*luc*). Expression of *cabABC* was then inferred by expression of *luc* and compared to known early (*comCDE*) and late (*sigX1* and *sigX2*) competence regulators. As expected, we observed that competence was not induced under acidic pH (Fig. S2), unless exogenous cognate competence-stimulating peptide (CSP_G22_) was supplied (Fig. S3A). Upon CSP_G22_ treatment, *cabABC* expression followed a similar temporal pattern to *comCDE*, *sigX1* and *sigX2*, peaking at approximately 70 min post-induction (Fig. S3B). Thus, based on expression kinetics alone, early and late competence genes could not be distinguished in this assay. This response was specific to CSP, as exogenous addition of cognate bacteriocin-like peptide pheromones (BlpC_1_ and BlpC_4_), encoded elsewhere in the *S. mitis* G22 genome (30), failed to induce *cabABC* expression (Fig. S4).

Deletion of the early regulator *comE* abolished *cabABC* expression, a result expected for both early-and late-regulated genes (Fig. 2B). Individual deletion of *sigX1* or *sigX2* partially reduced expression of *cabABC*, while deletion of both resulted in complete loss of *cabABC* expression, consistent with *cab* being expressed during the late phase.

Although multiple SigX boxes have been observed in *cab* promoters previously (16, 27), their functional relevance has not been tested. To assess this, we compared expression driven by the full-length *cabABC* promoter (containing all five SigX boxes) to a truncated version containing only the most proximal SigX box. Deletion of four out of five SigX boxes reduced *cabABC* expression (Fig. 2C), indicating that tandem SigX binding sites enhance *cab* expression. In the *S. mitis* collection, all *cab* loci harbored at least two SigX boxes, with one strain containing up to 38 boxes (Fig. 2D and Table S1). These boxes were often spread throughout *cab* loci, and larger loci were associated with higher number of SigX boxes, suggesting accumulation of boxes throughout these sequences.

Together, these results demonstrate that *cab* bacteriocins are specifically expressed during the late phase of competence, and that the presence of multiple SigX binding sites likely amplifies their expression.

### Expression of *S. mitis cabABC* is induced by pneumococcal CSP-1

Mitis group streptococci naturally produce and recognize different CSP variants, known as pherotypes (31). To investigate the specificity of *cab* expression, synthetic CSP variants from closely related strains were tested for their ability to activate *cabABC*. CSP variants from three other *S. mitis*, one *S. oralis*, and one *S. pneumoniae* strain failed to induce *cab* expression in *S. mitis* G22 (Fig. 3A). By contrast, the most prevalent *S. pneumoniae* pheromone, CSP-1 (32), efficiently induced *cab* to the same levels as native CSP_G22_.

**Fig. 3.**
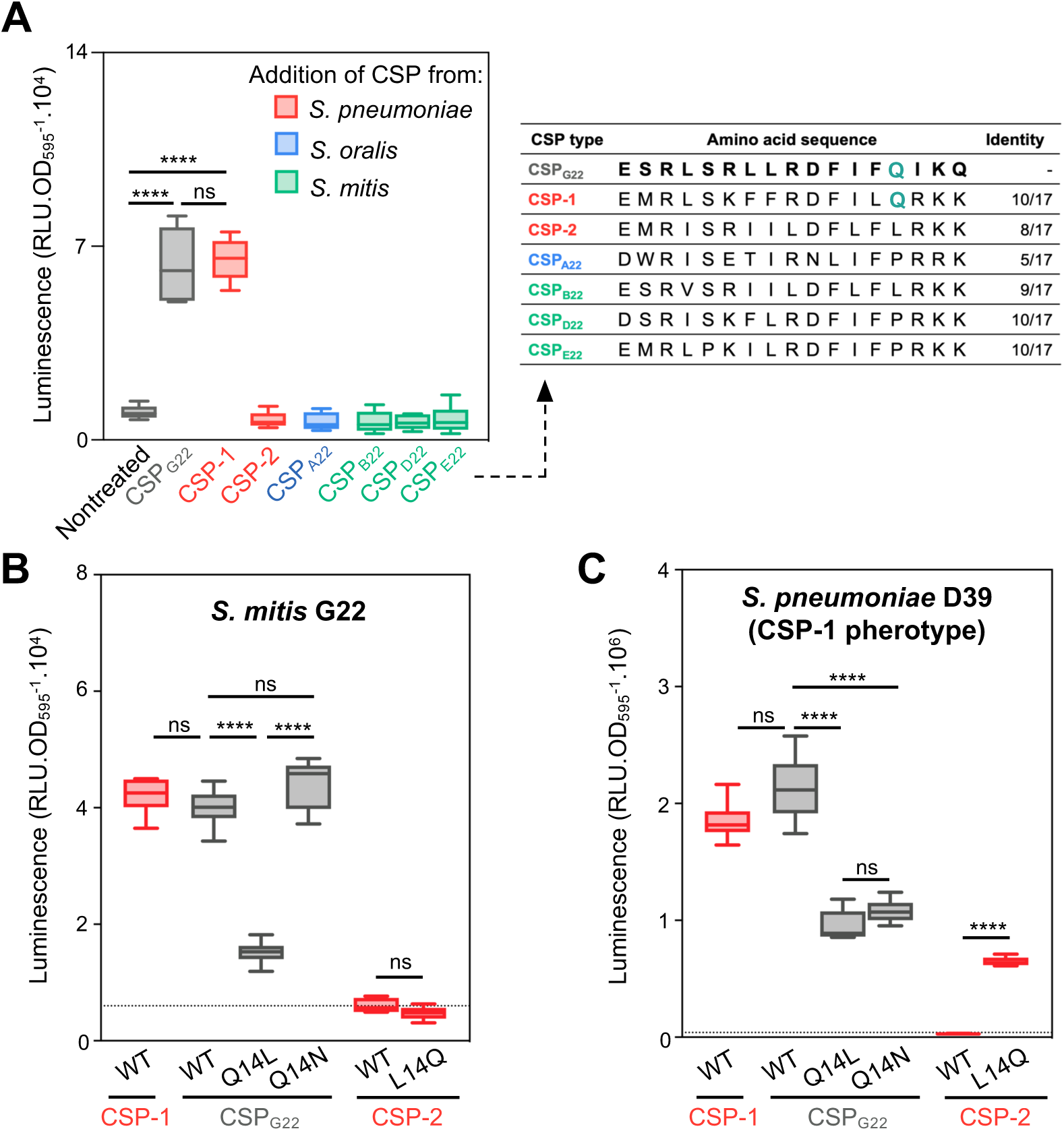
*S. mitis* and *S. pneumoniae* crosstalk through production of distinct CSP pheromones. (A) Effect of CSP pheromones of close streptococci on *cabABC* expression in *S. mitis* G22 and alignment of tested CSP variants (purple – Q14 amino acid residue, present only in inducing peptides). *S. mitis* G22 P*cabABC*-*luc* was pre-grown in acidic medium, to which exogenous CSP was added. (B) Effect of CSP with substitutions in residue 14 in the activation of *cabABC* expression in *S. mitis* G22. (C) Effect of CSP with substitutions in residue 14 in the activation of competence in *S. pneumoniae* of pherotype CSP-1 (P*comCDE*-*luc*). Dotted lines represent average luminescence value for nontreated controls. Experiments were repeated three times, independently. Unpaired *t*-tests were performed to compare *cab* expression. **** *P* < 0.0001. ns not significant.

Comparison of the amino acid sequences of CSP_G22_ and the other CSP variants revealed a single amino acid residue, Q14 (glutamine), that may explain the differences in induction. While both CSP_G22_ and CSP-1 contain Q14, a polar residue, the non-inducing variants contain either a L14 (leucine) or a P14 (proline), both nonpolar. To investigate the importance of Q14 and polarity in competence induction, CSP_G22_ variants with single amino acid substitutions at position 14 were tested. Substituting Q14 with L significantly reduced *cab* expression, whereas replacing it with another polar residue (N, asparagine) restored expression (Fig. 3B). This suggests that polarity of amino acid 14 is important for CSP recognition by *S. mitis* G22. However, introducing Q14 into a non-inducing peptide (CSP-2; L14Q) was not sufficient to convert it into a functional inducer, indicating that additional residues beyond Q14 are required for efficient recognition by the membrane-bound histidine kinase ComD of *S. mitis* G22.

To assess whether *S. pneumoniae* can also respond to CSP_G22_, the *comCDE* promoter from *S. pneumoniae* D39, a strain that produces and recognizes CSP-1, was transcriptionally fused to a *luc* reporter and integrated in its chromosome. Competence in D39 was efficiently induced by both CSP-1 and CSP_G22_, confirming that *S. mitis* G22 and *S. pneumoniae* D39 can cross-communicate through distinct CSP peptides (Fig. 3C). Similar to G22, *S. pneumoniae* D39 was also affected by mutations at position 14: Q14L (polar to nonpolar) reduced induction. In contrast to *S. mitis* G22, the substitution Q14N (polar with another polar amino acid), failed to restore it. Notably, the substitution L14Q into CSP-2 (the second most frequent CSP in pneumococci) enabled competence induction.

These findings highligh the importance of Q14 in CSP recognition of pneumococci of the CSP-1 pherotype.

In the 253 *S. mitis* genomes analyzed, 2.8% encode CSP_G22_ (Table S1) indicating that more *S. mitis* strains can induce competence in pneumococci of pherotype CSP-1.

These results reveal that cross-communications via CSP is possible between *S. mitis* and *S. pneumoniae*, and that residue 14 plays a critical, but context-dependent, role in pheromone recognition.

### CabAB mediates inhibition of *S. pneumoniae* in a contact-dependent manner

Given that *S. mitis* G22 and *S. pneumoniae* D39 can cross-communicate through CSP, we hypothesized both species would engage in competence-induced predation. Supporting this, *S. mitis* G22 harbors the orphan immunity gene *cibC* (but not *cibAB*), suggesting it may be resistant to pneumococcal CibAB attack (Table S1).

To investigate the specific role of the competence associated bacteriocins CabAB in *S. pneumoniae* inhibition, we first deleted two bacteriocin clusters (*blp4* and *blp1*) with known anti-pneumococcal activity (27), from the genome of *S. mitis* G22, as their activity could obscure CabAB-mediated effects. All experiments in this section were, thus, performed using *S. mitis* G22 Δ1*blp4*Δ1*blp1* and its derivatives as described below.

We grew dual-species biofilms of *S. mitis* G22 Δ1*blp4*Δ1*blp1* and *S. pneumoniae* D39 under competence-permissive conditions. After 24 h, both species persisted in the biofilm (Fig. 4A and Fig. S5). Deletion of *cab* from *S. mitis* led to increased pneumococcal loads, while constitutive expression of *cab* via plasmid p(*cab*^+^) (overexpressor mutant contains the native *cab* copy and carries an additional constitutively expressed copy in plasmid p(*cab*^+^)) reduced them, indicating that the presence and expression levels of *cab* modulate competitive outcomes.

**Fig. 4.**
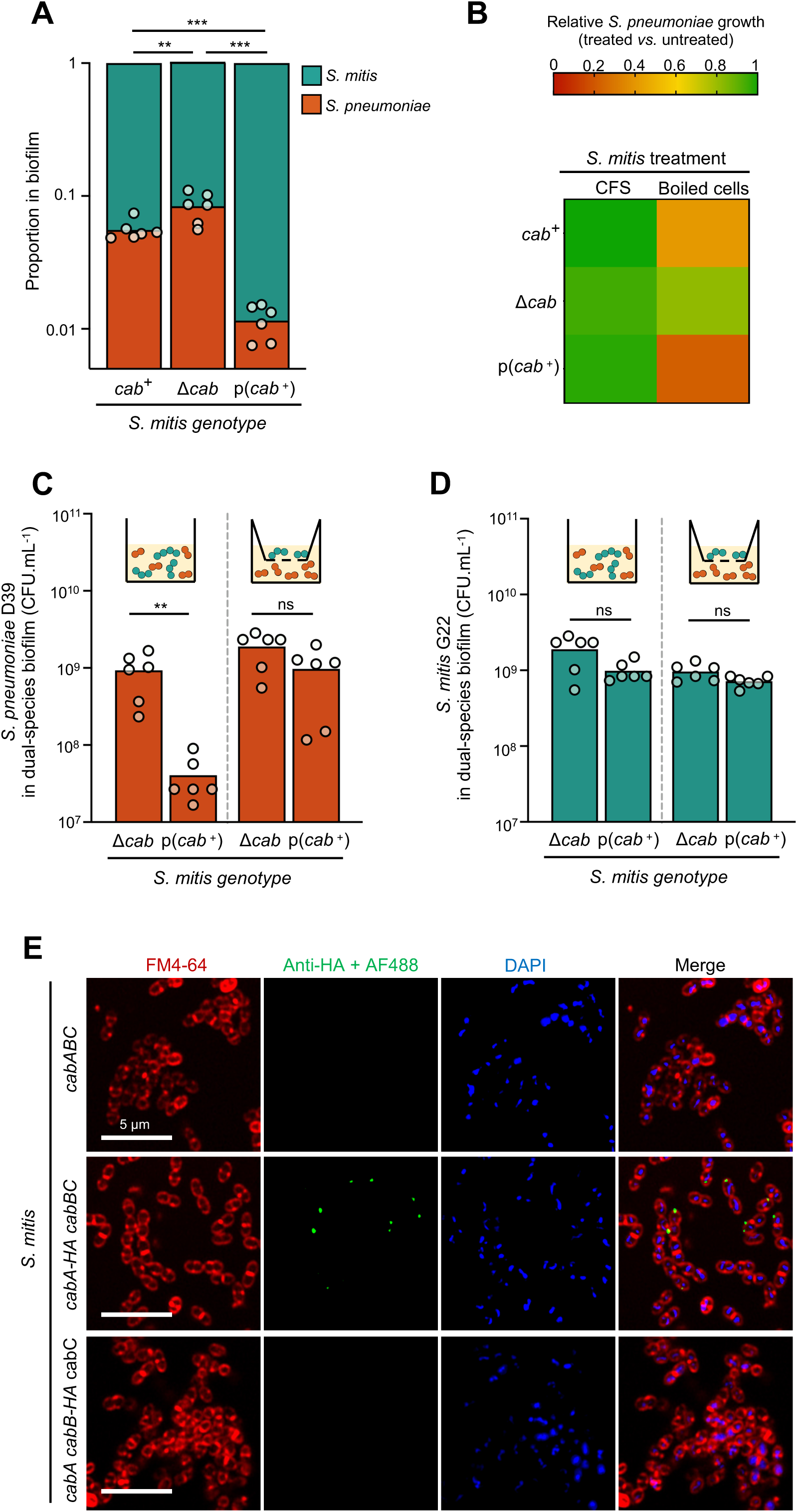
*S. mitis* bacteriocins CabAB mediate contact-dependent growth inhibition of *S. pneumoniae*. Experiments were done using *S. mitis* G22 Δ1*blp4*Δ1*blp1* and *S. pneumoniae* D39. The bacteriocin clusters *blp4* and *blp1* were first deleted from the genome of *S. mitis* G22 as they have anti-pneumococcal activity (27), which could obscure CabAB-mediated effects. (A) Proportion of *S. mitis* and *S. pneumoniae* in dual-species biofilm grown for 24 h in competence-permissive conditions in 24-well plates. *S. mitis* mutant p(*cab*^+^) contains the native *cabABC* copy, and an additional copy under constitutive regulation (promoter *p23*) in a multicopy plasmid. (B) *S. mitis* was grown in competence-permissive conditions. Cell-free supernatants (CFS) were recovered and cell pellets were boiled. CFS and heat-killed cells were added to *S. pneumoniae* cultures and growth was monitored. (C) Bacterial loads of *S. pneumoniae* grown in the presence of *S. mitis* in direct contact (left) or separated with a 0.4 μm pore transwell membrane (right). Biofilms were grown in 12-well plates. As the surface area available for cell attachment is higher in 12-well plates than in 24-well plates, we hypothesize that this contributed to the observed differences in maximum cell counts for D39. (D) Corresponding bacterial loads of *S. mitis* G22 in experiment (C). (E) Representative images of CabA-HA and CabB-HA immunodetection on the surface of live competent *S. mitis* cells containing the native *cabABC* (negative control), *cabA-HA cabBC* or *cabA cabB-HA cabC* genotypes. *S. mitis* cultures were induced with CSP_G22_, incubated with a primary anti-HA rabbit antibody, and then with an Alexa Fluor 488-conjugated goat anti-rabbit secondary antibody. Cells were stained with FM4-64 (membrane dye) and DAPI (nucleic acid dye). At least ten independent fields were detected per experiment/condition. Experiments were repeated at least three times, independently. The ratio paired *t*-test was used to compare bacterial proportions in dual-species biofilms. The two-tailed unpaired Mann-Whitney U test was used to compare bacterial loads. ** *P* < 0.01. *** *P* < 0.001. ns not significant.

Bacteriocins are typically secreted to the extracellular environment (33). To test whether CabAB exerts its activity as a secreted factor, we harvested cell-free supernatants (CFS) from *S. mitis* strains and added them to pneumococcal cultures. CFS of *S. mitis* natively or overexpressing *cab* did not inhibit *S. pneumoniae* growth (Fig. 4B), suggesting CabAB is not released into the medium. This aligns with recent reports of bacteriocins, such as Listeriolysin S and contact-dependent inhibition by glycine zipper proteins CdzCD (34, 35), that remain attached to the producer cell surface. In these cases, inhibitory activity occurs via direct contact between producer and target cells.

To test whether CabAB could remain cell-associated, we heat-killed *S. mitis* strains (as bacteriocins are typically resistant to high temperature treatments) and exposed them to *S. pneumoniae*. Heat-killed *cab*^+^ and p(*cab*^+^) *S. mitis* extracts significantly inhibited pneumococcal growth, while extracts from Δ*cab S. mitis* did not (Fig. 4B), supporting the idea that CabAB is retained in producer cells.

We then tested whether direct contact between *S. mitis* and *S. pneumoniae* was required for inhibition. Dual-species biofilms were grown either with or without a 0.4 µm transwell membrane separating both species. When the strains were in direct contact with each other, *cab* expression was associated with pneumococcal inhibition, whereas no inhibition was observed when both species were separated (Fig. 4C). These effects were not due to differences in *S. mitis* growth, as bacterial loads were consistent across conditions and genotypes (Fig. 4D).

We then sought to detect CabA and CabB on the surface of competent *S. mitis* cells. A mutant lacking the native *cab* locus was used, into which we ectopically integrated cassettes containing haemagglutinin-tagged (HA) CabA (*cabA-HA cabBC*) or CabB (*cabA cabB-HA cabC*), under native regulation. Immunofluorescence microscopy of live *S. mitis* expressing *cabA-HA* using anti-HA antibodies revealed discrete clusters of CabA on the cell surface (Fig. 4E). Of note, CabA was not detectable in all cells. This may be due to several factors, such as CabA being occluded by the cell wall or competence development being asynchronous, with different subpopulations of *S. mitis* becoming competent at slightly different times. No signal was detected in strains lacking the HA epitope, confirming specificity. Interestingly, no signal was observed in the *cabB-HA* strain, suggesting that CabB may be less surface-exposed, that the N-terminal HA tag was not accessible to antibody binding, or that tagging interfered with localization.

Together, these results demonstrate that *S. mitis* inhibits *S. pneumoniae* via CabAB in a contact-dependent manner.

### CabAB is a two-peptide bacteriocin with LytA-dependent bactericidal activity

CibAB has previously been shown to be a two-peptide bacteriocin, meaning its inhibitory activity occurs only when both peptide components are present (8). To test if CabAB also functions as a two-peptide bacteriocin, synthetic CabA and CabB were tested individually and in combination against *S. pneumoniae* D39. No effect on pneumococcal growth was observed upon addition of DMSO (the solvent used to resuspend the peptides), CabA alone, or CabB alone (Fig. 5A). In contrast, a single treatment with equimolar amounts of CabA and CabB delayed *S. pneumoniae* growth for ∼12 h, indicating that both peptides are required for activity. This is consistent with the presence of multiple GxxxG or GxxxG-like motifs in both bacteriocins (Fig. 5B), known to facilitate and stabilize two-component bacteriocins (36–38). We also confirmed that CabAB can cross the membrane used in the transwell assays, leading to a decrease in *S. pneumoniae* loads when added to opposite compartments (Fig. S6).

**Fig. 5.**
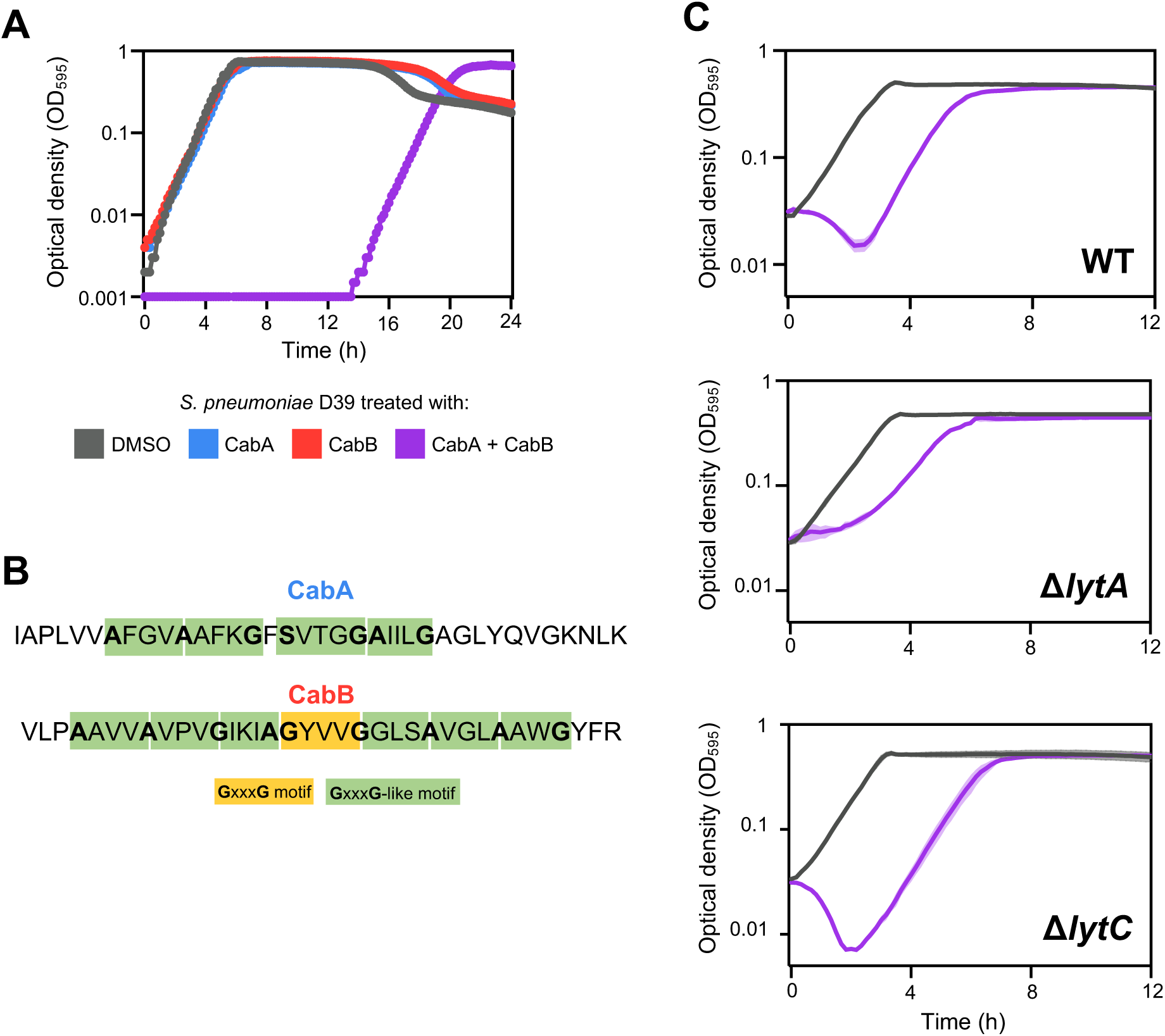
CabAB is a two-peptide bacteriocin with LytA-dependent bactericidal activity against *S. pneumoniae.* (A) *S. pneumoniae* was treated with synthetic CabA, CabB, an equimolar mixture of CabA and CabB (0.975 µM), or with DMSO (solvent where peptides were resuspended). Growth was recorded for 24 h. (B) Presence of GxxxG and GxxxG-like amino acid motifs in CabA and CabB sequences. (C) Exponential cultures of wild-type and autolysin-deficient *S. pneumoniae* were treated with DMSO (grey) or synthetic CabAB (purple). Growth was recorded for 24 h. For easier visualization of lysis (decrease in optical density), the first 12 h of growth is represented. Growth curves were repeated three times independently, and one representative experiment with two technical replicates per condition is shown.

We next investigated whether CabAB exerts bactericidal activity. Exponentially growing *S. pneumoniae* D39 cultures were treated with CabAB or DMSO. CabAB-treated cultures showed evident decrease in optical density, indicating lysis, while DMSO-treated controls did not (Fig. 5C). Lysis induced by the related bacteriocin CibAB has been shown to depend on the action of the pneumococcal autolysins LytA and LytC (8). Here, deletion of *lytC* from D39 had no effect on CabAB-mediated lysis, whereas a Δ*lytA* mutant exhibited delayed growth but no decrease in optical density, indicating that LytA plays a key role in CabAB-induced lysis. This does not exclude, however, the possibility that limited lysis occurs concomitantly with growth.

Together, these findings show that *S. mitis* CabAB is a two-peptide bacteriocin with bactericidal activity against *S. pneumoniae*, and that its lytic activity depends on the target cell LytA autolysin.

### CabAB affects membrane homeostasis in *S. pneumoniae*

We sought to shed light on how CabAB affects pneumococcal physiology. To this end, *S. pneumoniae* D39 cells were treated with CabAB or DMSO and imaged by scanning electron microscopy (SEM).

While DMSO-treated cultures displayed normal morphology, CabAB-treated cells appeared to be encased in an extracellular matrix (Fig. 6A), suggesting leakage of intracellular material.

**Fig. 6.**
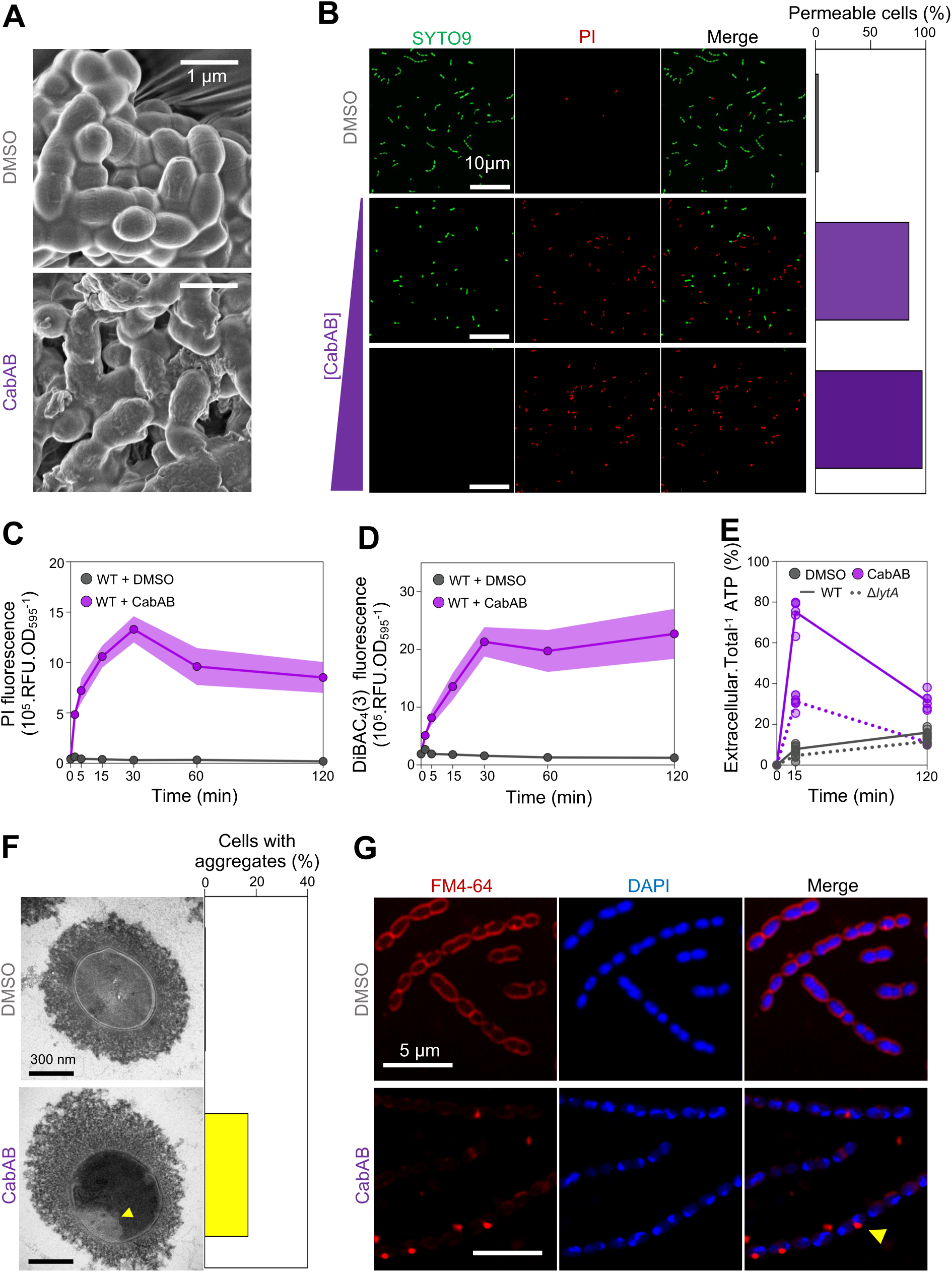
CabAB leads to permeabilization and depolarization of *S. pneumoniae* membranes, leading to metabolite leakage. (A) *S. pneumoniae* treated with DMSO or CabAB and imaged by scanning electron microscopy (SEM). (B) Quantification of membrane-permeable *S. pneumoniae* (stained with propidium iodide (PI)) upon treatment with DMSO (560 cells) or increasing concentrations of CabAB (0.975 µM – 481 cells; or 1.95 µM – 397 cells). (C-D) Quantification of populational membrane permeability (C) and depolarization (D) upon DMSO or CabAB treatment at several timepoints. (E) Proportion of extracellular *vs.* total ATP in wild-type and Δ*lytA S. pneumoniae* cultures upon DMSO or CabAB treatment. (F) Representative transmission electron microscopy (TEM) image and quantification of intracellular aggregates in DMSO-(n = 118 cells) or CabAB-treated *S. pneumoniae* (n = 138 cells). (G) Labeling of large intracellular aggregates by membrane dye FM4-64 in CabAB-treated *S. pneumoniae*. SEM and TEM were done once. Other data was taken from at least three independent experiments.

To test if membrane permeability was compromised, cells were stained with propidium iodide (PI), a DNA-binding dye that only enters cells with compromised membranes. In DMSO-treated cultures, only 3% were PI-positive, while CabAB-treated cultures contained 85% PI-positive cells (Fig. 6B). Doubling the concentration of CabAB further increased this proportion to 97%, indicating this phenotype is concentration dependent. Deletion of *lytA* had no effect on these phenotypes, demonstrating that membrane permeabilization occurs independently of autolysin-mediated lysis (Fig. S7).

Time series measurements showed that PI signal intensity peaked at 30 min post CabAB exposure, still being detected after 120 min (Fig. 6C). We also assessed pneumococcal membrane depolarization using the fluorescent dye DiBAC_4_(3), which enters cells only when their membranes are depolarized. Compared to DMSO, CabAB treatment led to a rapid and sustained increase in DiBAC_4_(3) signal (Fig. 6D), indicating membrane depolarization.

Since some characterized two-peptide bacteriocins act by forming pores (39–42), we tested whether CabAB treatment causes efflux of intracellular components from target cells. In wild-type *S. pneumoniae* treated with CabAB, 80% of total ATP was detected in the supernatant (Fig. 6E), in agreement with our previous observation of significant cell lysis (Fig. 5C). Interestingly, in CabAB-treated Δ1*lytA S. pneumoniae* cells, no decrease in optical density was observed over the course of the experiment, yet 30% of ATP was still detected in the supernatant, suggesting that CabAB induces leakage independently of cell lysis.

We next visualized CabAB-induced damages using transmission electron microscopy (TEM). DMSO-treated cells appeared intact, whereas 17% of CabAB-treated cells contained large intracellular aggregates (Fig. 6F). Based on our prior observations, we hypothesized these aggregates would be membranous. Consistently, staining with the membrane dye FM4-64 revealed intracellular foci in CabAB-treated cells, but not in control cells (Fig. 6G), confirming the formation of membrane aggregates by CabAB.

Finally, lower-magnification fluorescent microscopy of CabAB-treated cultures stained with DAPI and FM4-64 revealed frequent clumps of cells encased in nucleic acid and membrane material, mirroring the SEM observations of surviving cells remaining embedded in an extracellular matrix (Fig. S8)

Altogether, these findings show that *S. mitis* CabAB disrupts membrane homeostasis in *S. pneumoniae* by permeabilizing and depolarizing membranes, leading to release of its intracellular content.

### CabAB facilitates the acquisition of pneumococcal DNA by *S. mitis* in biofilms

As previously shown, CabAB promotes release of DNA from *S. pneumoniae* cultures. Since *cab* expression is synchronized with the competence period, we hypothesized that CabAB could enhance DNA acquisition and HGT within biofilms. To test this, we grew dual-species biofilms under conditions designed to study HGT (Fig. 7A), where: (i) DNA transfer is unidirectional, since the DNA donor strain (*S. pneumoniae* D39) carries a deletion of the major competence pilus subunit gene, *comGC,* rendering it nontransformable; (ii) the *S. pneumoniae* DNA donor harbors chloramphenicol (Chl^R^) and tetracycline (Tet^R^) resistance cassettes integrated at opposite regions of the chromosome with high sequence homology (89-98%) between DNA donor and recipient (Fig. S9); (iii) the DNA recipients (*S. mitis* G22, wt and variants) are kanamycin resistant, allowing for discrimination between non-transformed (Kan^R^) and transformed *S. mitis* (Kan^R^ Chl^R^, Kan^R^ Tet^R^, and Kan^R^ Chl^R^ Tet^R^); (iv) biofilms are pre-grown under acidic pH for 24 h to promote biofilm establishment without inducing competence; and (v) CSP_G22_ is added after 24 h to transiently induce competence. A noncompetent *S. mitis* Δ*comE* mutant was used as a negative control, as it is unable to acquire Chl^R^ and/or Tet^R^ by transformation. These experiments were conducted using the *S. mitis* G22 Δ1*blp4* Δ1*blp1* background.

**Fig. 7.**
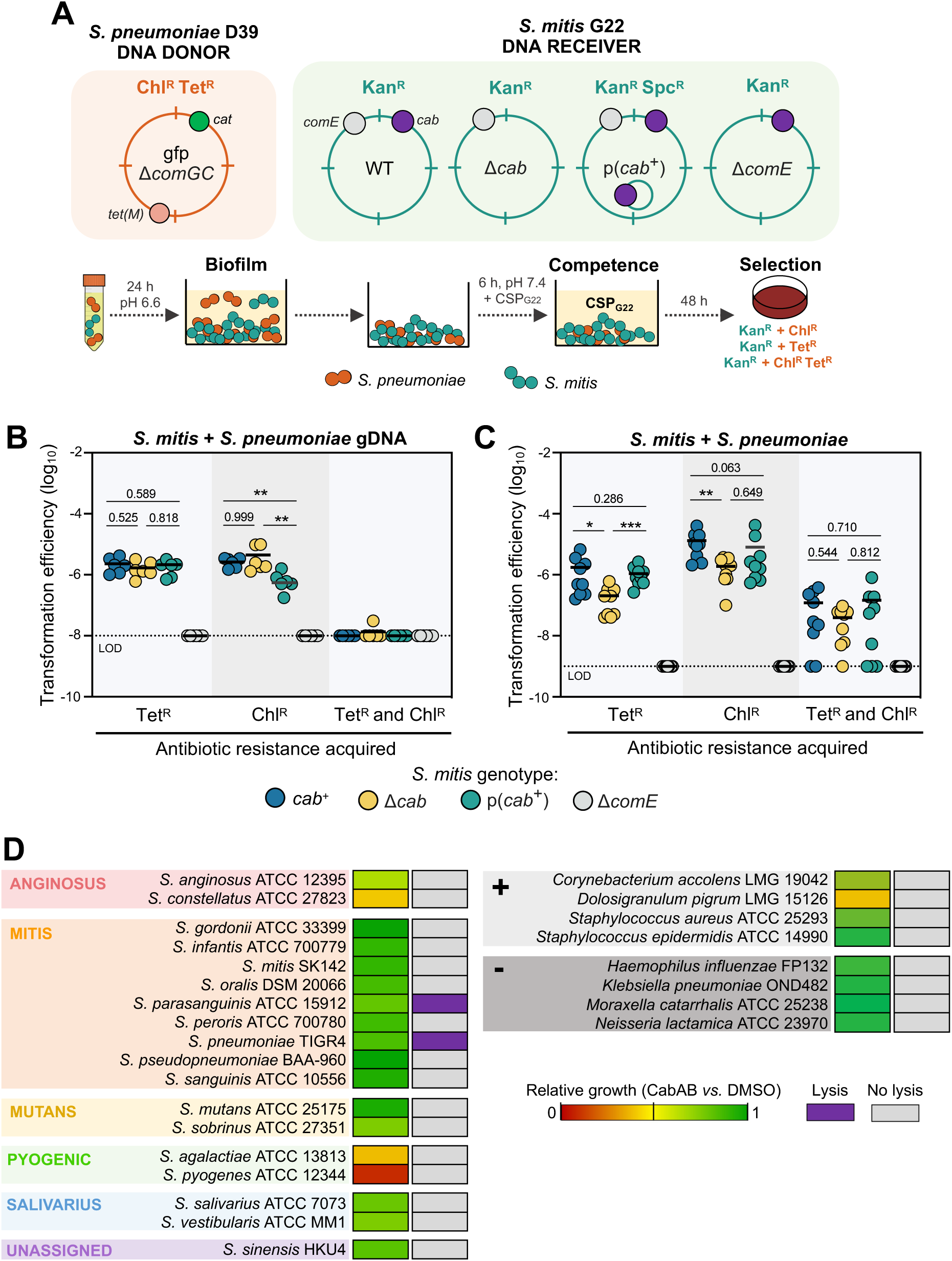
Expression of *cabAB* facilitates horizontal gene transfer in dual-species biofilms. (A) Experimental setup to study gene transfer in dual-species biofilms. The *S. pneumoniae* DNA donor is nontransformable due to a *comGC* deletion, and contains chloramphenicol and tetracycline resistance cassettes at opposite ends of the chromosome (Chl^R^ and Tet^R^); the *S. mitis* DNA receiver is labeled with a kanamycin resistance cassette (Kan^R^); biofilms were pre-grown at a starting 1:1 ratio for 24 h in acidic medium to prevent natural competence induction; cells were aspirated and fresh medium containing CSP_G22_ was added to biofilms to transiently induce competence; after 6 h of incubation, cells were plated; *S. mitis* transformants were recovered and counted by selection in plates with kanamycin and combinations of chloramphenicol and/or tetracycline. (B-C) Acquisition of Chl^R^ and/or Tet^R^ by *S. mitis* in biofilms containing purified donor gDNA (B) or the strain itself (C). (D) Growth inhibition and lysis of 26 bacterial species when treated with 0.975 µM CabAB *vs.* DMSO during their exponential phase. Growth was recorded for 24 h. Lysis is shown when a decrease from the initial culture optical density was observed in all replicates. Experiments were repeated three times, independently. Transformation efficiencies were compared using unpaired *t*-tests. LOD limit of detection. * *P* < 0.05. ** *P* < 0.01.

To rule out the possibility that CabAB affects transformability rather than DNA availability, we first replaced the donor strain with purified gDNA. Under these conditions, both *cab*^+^ and Δ1*cab S. mitis* acquired resistance at similar frequencies (Fig. 7B), indicating that native *cab* expression does not impact transformability. In contrast, overexpression of *cab* reduced transformation efficiency for one antibiotic resistance, possibly due to the fitness burden associated with carrying p(*cab*^+^).

In dual-species biofilms, *cab*^+^ *S. mitis* efficiently acquired Chl^R^ and Tet^R^ from the DNA donor, at a significantly higher frequency than the Δ1*cab* mutant (Fig. 7C), indicating that CabAB contributes to DNA acquisition. In contrast, overexpression of CabAB did not enhance transformation efficiency, suggesting that a precise *cab* regulation might be necessary for optimal transformation. We extended these findings to an additional *S. mitis* strain, C22, which harbors a distinct *cab* bacteriocin repertoire (27). *S. mitis* C22 contains a *cab* cluster encoding two copies of the bacteriocin *cabD* (cluster 10), two copies of the bacteriocin *cabE* (cluster 26), and two putative immunity proteins, *cabF* and *cabG* (Fig. S10A). Wild-type C22 was able to acquire Chl^R^ and Tet^R^ from the *S. pneumoniae* DNA donor and transformation frequencies were also reduced in the Δ1*cab* mutant (Fig. S10B). Consistently, the proportion of *S. pneumoniae* in dual-species biofilms with *S. mitis* C22 was lower when co-cultured with C22 *cab*^+^ than with Δ1*cab* cells (Fig. S10C), showing that the cab locus of a different *S. mitis* strain also mediates inhibition of *S. pneumoniae* and facilitates genetic exchange.

We then sought to investigate the specificity of CabAB-induced lysis. Synthetic CabAB was tested against one representative strain of 26 bacterial species, including 18 streptococcal species, four additional non-streptococcal Gram-positive and four Gram-negative species. All selected species are human commensals or pathogens that can inhabit the oral and/or respiratory tract and could theoretically encounter *S. mitis* and be exposed to Cab bacteriocins in natural environments. Addition of 0.975 µM of CabAB to exponential cultures of these bacteria stalled the growth of some species (Fig. 7D and Fig. S11 and S12). The most sensitive to CabAB were the representative strains of Pyogenic Group streptococci (*S. pyogenes* and *S. agalactiae*). Gram-negative bacteria (*Haemophilus influenzae*, *Klebsiella pneumoniae*, *Moraxella catarrhalis*, and *Neisseria lactamica*) were largely unaffected. Despite growth inhibition in some species, lysis was only observed in two - *S. pneumoniae* and *S. parasanguinis* - suggesting that CabAB-mediated lysis has limited specificity.

Finally, we investigated whether membrane aggregate formation was specifically associated with species that undergo CabAB-mediated lysis. Aggregate formation was observed exclusively in *S. pneumoniae* and not in *S. parasanguinis*, *S. pyogenes* and *S. agalactiae* (Fig. S13). Notably, membrane aggregates were also absent in Δ1*lytA S. pneumoniae* cells, indicating that aggregate formation is specific to *S. pneumoniae* and that it requires LytA.

These results suggest that CabAB provides a competitive advantage by inhibiting the growth of some bacteria, but that it also enhances DNA release in biofilms by selectively lysing a few closely related species during competence, notably *S. pneumoniae*, thereby promoting gene exchange.

## Discussion

During competence, streptococci scavenge extracellular DNA. In *S. pneumoniae*, where competence-induced predation is best characterized, the coordinated action of CibAB, LytA, LytC, and the murein hydrolase CbpD leads to lysis of noncompetent siblings (8). This process promotes DNA release and facilitates HGT (12, 43, 44).

Although extensive genomic data suggests that *S. mitis*, a close relative of *S. pneumoniae*, frequently acquires *S. pneumoniae* DNA (18, 19, 21–24, 45), the molecular mechanisms underlying this exchange remained poorly understood. Understanding these mechanisms is relevant because capsule acquisition by *S. mitis* can complicate DNA-based pneumococcal serotyping. In addition, *S. mitis* has been associated with cases of infective endocarditis (23), highlighting the value of studying how it acquires potentially pathogenic traits.

Here, we investigated the role of *S. mitis* competence-associated bacteriocins, CabAB, in mediating interspecies interactions (Fig. 8). We show that CabAB is a two-peptide bacteriocin produced during the late phase of competence, when cells are primed for DNA uptake and recombination, capable of lysing *S. pneumoniae* in a LytA-dependent manner. Although LytA is constitutively expressed throughout growth, its concentration increases during competence, owing to a genomic architecture comprising three constitutive promoters and one competence-induced promoter (7, 46–48). In this context, it is tempting to speculate that by inducing *S. pneumoniae* competence via CSP crosstalk, and therefore increasing LytA concentration, some *S. mitis* could further sensitize *S. pneumoniae* to Cab bacteriocins. Nonetheless, CabAB-induced lysis appears to be independent of pherotype of target cells, as both *S. pneumoniae* D39 (CSP-1; used in most experiments) and TIGR4 (CSP-2; included in the panel of 26 bacterial species) were lysed. These pherotypes account for the vast majority (>95%) of pneumococcal strains (49–51).

**Fig. 8.**
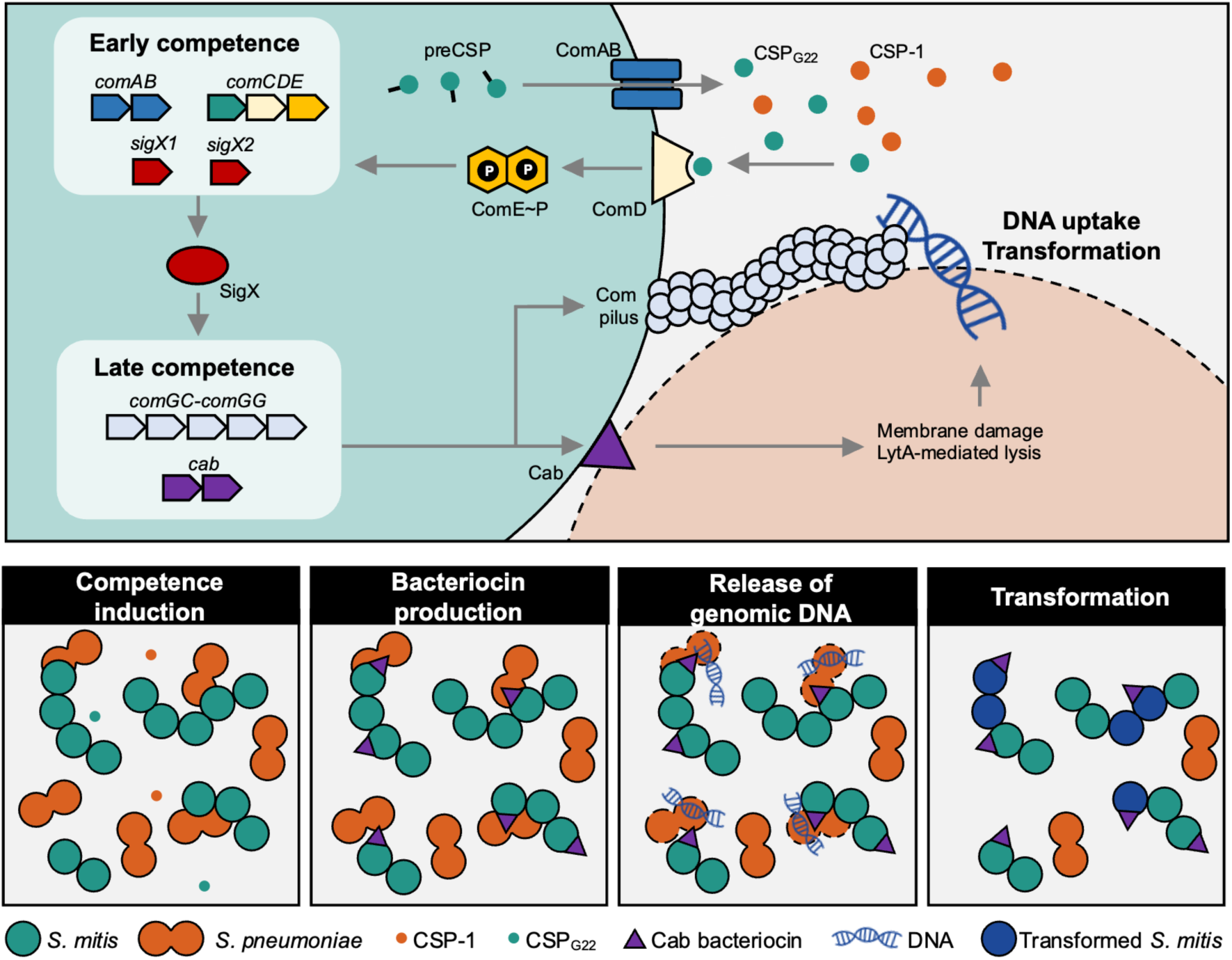
Model for the role of Cab bacteriocins in the acquisition of heterologous DNA. *S. mitis* produces the CSP precursor (preCSP) intracellularly, which is exported and processed by ComAB. Extracellular CSP(CSP_G22_), as well as other CSP variants such as pneumococcal CSP-1, may be recognized by the membrane-bound histidine kinase ComD, leading to phosphorylation of the response regulator ComE. Phosphorylated ComE (ComE∼P) activates expression of the early competence genes, establishing a positive feedback loop. This includes *sigX1 and sigX2*, which encode for the alternative sigma factor that drives expression of late competence genes. Among the late genes are the *cab* bacteriocins. Cab localize to the surface of producer cells. Surface-associated Cab mediate lysis of neighbouring *S. pneumoniae* cells in a cell contact-and LytA-dependent manner, resulting in the release of DNA. During the late competence phase, *S. mitis* cells also assemble a competence pilus responsible for the uptake of extracellular DNA, which can subsequently be integrated into the genome by homologous recombination.

Of the initial 332 *S. mitis* genomes analyzed, we were not able to retrieve the region upstream of *comAB* from a substantial fraction of strains (23.8%), as their assemblies showed contig breaks precisely at this locus. We propose this difficulty arises from the repetitive nature of the region, as indicated by the presence of numerous SigX boxes. The number of boxes scaled linearly with the size of *cab* loci, suggesting that SigX boxes have accumulated throughout these sequences. Repeats are known to both promote and result from genomic rearrangements (52, 53). They are often remnants of previous recombination events, with no known biological function (54). In this case, however, accumulation of SigX boxes may provide a selective advantage by enhancing bacteriocin expression, which could explain why they are enriched in this region. Consistent with this, the extensive genetic rearrangements observed across multiple streptococcal species at this locus also suggests this is a highly recombinogenic region (25, 26, 55). To the best of our knowledge, the *cab* locus appears to be the only competence-induced locus in *S. mitis* whose promoters contain multiple SigX boxes; other competence-induced genes typically harbor a single box (16).

Microscopic analysis revealed that CabAB-treated pneumococci often contain large membrane-derived intracellular aggregates. Whether these structures are part of CabAB’s mechanism of action or represent a cellular stress response remains unclear. Intracellular vesicle formation has been shown to be essential for the lytic activity of bacteriocin A37 (56). Consistently, we have observed that deletion of LytA, required for CabAB-mediated lysis in *S. pneumoniae*, blocks formation of intracellular aggregates.

We further demonstrate that CabAB-mediated inhibition of *S. pneumoniae* requires direct contact between producer and target cells. Contact-dependent bacteriocin delivery has been described in *Caulobacter crescentus* and *Listeria monocytogenes* (34, 35). In *L. monocytogenes*, bacteriocins were only detected on the surface of producer cells following cell wall digestion. In contrast, we detected CabAB on live *S. mitis* with intact cell walls, suggesting a distinct delivery mechanism. Although CabAB activity strictly required both peptides, only CabA could be detected on the surface of competent *S. mitis*. CabB was not detectable by immunofluorescence, which could reflect limited surface exposure, inaccessibility of the HA tag, or altered localization caused by C-terminal tagging. A similar pattern of differential detectability has been reported in the CdzC/D contact-dependent bacteriocin system in *Caulobacter crescentus*: both CdzC and CdzD are cell-associated and required for activity, but only CdzC is clearly visualized in surface aggregates, whereas CdzD is not direcly observed on the surface (35). These observations suggest that, in two-peptide contact-dependent killing systems, one peptide may be more readily surface-exposed. Contact-dependent killing during competence is not unique to streptococci, despite differences in killing factors. *Vibrio cholerae* produces a competence-induced type VI secretion system that kills nonimmune cells, releasing DNA that becomes accessible for HGT (57). Contact dependence may be particularly advantageous in biofilm contexts, a niche where extracellular DNA could be subject to rapid degradation nucleases (58). Proximity within multispecies biofilms could enhance transformation efficiency by increasing the local DNA concentration, given that DNA abundance and size influence recombination success (59). Notably, biofilm conditions and cell-to-cell contact have been shown to promote both the frequency and size of recombination events in pneumococci, respectively (60, 61). Notably, the frequency of double transformants was higher under cocultivation conditions (*S. mitis* and *S. pneumoniae*) than when genomic DNA was added to *S. mitis* biofilms. In coculture, DNA is likely released continuously over the course of incubation, increasing its local concentration and availability, which may favor sequencial uptake and recombination events. In contrast, purified gDNA was added only once and could be subject to degradation by extracellular nucleases, thereby reducing the probability of independent recombination events. Strikingly, among 26 tested species, only *S. pneumoniae* and *S. parasanguinis* were lysed by CabAB, suggesting a narrow-spectrum bactericidal activity. Whether this specificity is unique to CabAB or shared among other Cab bacteriocins remains to be determined. Nonetheless, the ability to selectively target phylogenetically related but distinct species suggests a potential trade-off between maximizing recombination efficiency and accessing a broader gene pool. While high sequence identity between donor and recipient promotes recombination (62), restricting lysis to closely related species could expand the range of accessible alleles beyond conspecifics (*i.e.* beyond members of the same species). Thus, the large diversity of *cab* sequences in *S. mitis*, in contrast to *S. pneumoniae*, which typically harbors only *cibAB*, may represent one of several evolutionary forces contributing to the exceptional genetic diversity observed in *S. mitis* (23, 63).

Importantly, CabAB also strongly inhibited the growth of other streptococcal pathogens, including *S. pyogenes* and *S. agalactiae*, both of which are listed as priority pathogens by the World Health Organization (30). This potent inhibitory activity suggests that the Cab arsenal carried by *S. mitis* could be a promising source of antimicrobial agents against important clinical pathogens as recently suggested by us (27).

In summary, our findings uncover a contact-dependent mechanism of bacteriocin delivery in *S. mitis* that selectively targets closely related streptococci including *S. pneumoniae*. This process promotes HGT and likely contributed to competitive fitness and evolutionary plasticity of streptococcal populations.

## Materials and Methods

### Bacterial strains and culture conditions

Bacterial strains, oligonucleotides, plasmids and peptides used in the study are listed in Table S2. Streptococci were routinely grown in C+Y_YB_ (64) (pH 7.4, unless otherwise specified) and plated on tryptic soy agar with 5% defibrinated sheep blood (blood agar, BA). When appropriate, media was supplemented with antibiotics at the following concentrations: 150 µg.mL^-1^ kanamycin, 200 µg.mL^-1^ spectinomycin, 4 µg.mL^-1^ chloramphenicol, or 4 µg.mL^-1^ tetracycline.

*Staphylococcus aureus, Staphylococcus epidermidis* and *Moraxella catarrhalis* were grown in tryptic soy broth (TSB); *Corynebacterium accolens* was grown in TSB with 0.5% Tween 80; *Klebsiella pneumoniae* and *Escherichia coli* were grown in lysogeny broth; and *Neisseria lactamica*, *Haemophilus influenzae* and *Dolosigranulum pigrum* were grown in C+Y_YB_ with 20 µg.mL^-1^ hemin and 20 µg.mL^-1^ NAD. For propagation of plasmids in *E. coli*, media was supplemented with 100 µg.mL^-1^ kanamycin or 100 µg.mL^-1^ spectinomycin, as required.

### Identification of *cab* bacteriocins and *cibABC* in *S*. *mitis*

Publicly available *S. mitis* genomes (n = 322) were downloaded from PubMLST. An additional 10 genomes from reference or commonly used *S. mitis* strains were included, resulting in a total of 332 genomes (Table S1). To detect *cab* clusters, nucleotide sequences of *plsX*-*acpP* and *comAB* (from *S. mitis* B6; GenBank Accession GCA_000027165.1), which flank the *cab* locus, were aligned against all genomes using pairwise sequence alignment (Biopython v1.81; match = 2; mismatch =-1; gap open =-5; gap extension =-0.5; minimum overall sequence identity ≥ 90%). Genomes in which *plsX*-*acpP* and *comAB* were absent or located on separate contigs were excluded from further analysis. This region was identified in 253 genomes.

To identify mature Cab bacteriocin sequences, extracted sequences were scanned for open reading frames (ORFs) of 100-300 nucleotides, beginning with an ATG start codon and ending with a TAA, TAG or TGA stop codon. Only ORFs whose translated sequences contained a (M/L/V)XXXXGG double-glycine cleavage motif were retained. Mature Cab bacteriocins were obtained by removing the N-terminal signal sequence upstream of the cleavage site. In this work, we defined *cab* bacteriocins as all non-*cib* bacteriocin genes that were found in this particular location. To explore diversity of Cab bacteriocins, sequences were converted into amino acid frequency vectors. Pairwise dissimilarities between sequences were calculated using the Bray-Curtis metric (ScyPy), followed by agglomerative hierarchical clustering (scikit-learn). To determine the optimal number of clusters, silhouette scores (scikit-learn) were computed for cluster numbers ranging from 2 to 50. The number of clusters yielding the highest score was selected as optimal.

To assess the prevalence of *cibABC* in *S. mitis*, the nucleotide sequence of *cibABC* (from *S. pneumoniae* D39, GenBank Accession NC_008533.2) was aligned against all *S. mitis* genomes using NCBI BLASTn. Genomes with no detectable hit were considered *cibABC*-negative. For genomes with partial matches (<100% query coverage), the locus was manually inspected to verify the presence or absence of *cibA*, *cibB*, and *cibC* individually – a common occurrence in strains harboring *cibC* but not *cibAB*.

To find SigX boxes, the consensus sequence (65) was searched within *cab* loci using Find Individual Motif Occurrences (FIMO) v5.5.8 (66). Hits with *p*-value <0.001 were considered true SigX boxes. The detection of Cab bacteriocins and putative SigX boxes was benchmarked against six *S. mitis* strains with complete genomes and manually curated *cab* loci (27, 67).

### Phylogeny of *S. mitis*

Core genome single-nucleotide polymorphisms (SNPs) were called using Snippy v4.6.0 (https://github.com/tseemann/snippy) by aligning *S. mitis* genomes to the *S. mitis* B6 reference genome. The resulting alignment was used to construct a maximum-likelihood phylogenetic tree with FastTree v2.2 (68). The phylogeny was visualized and annotated using the Interactive Tree of Life (iTOL) platform (69).

### Synthetic peptides

Peptides were produced by solid-phase peptide synthesis with >95% purity (NZYTech) (Table S2) (27, 49–51). CSP and BlpC_1_ pheromones were resuspended in sterile ddH_2_O; BlpC_4_ was resuspended in 10% acetic acid; and CabA and CabB were resuspended in DMSO. Peptides were serially diluted in sterile ddH_2_O and stored at-20°C.

### Transformation

For transformation of *S. mitis* and *S. pneumoniae*, cells were grown to an OD_600_ of 0.5, diluted 1:100 in pre-warmed C+Y_YB_ and grown until an OD_600_ of 0.04-0.1. Cells were treated with 1 µM of cognate CSP and transformed with 0.4 µg linear DNA, 1 µg plasmid DNA, or 10 µg chromosomal DNA. Transformants were recovered for 4 h and plated in selective medium. Linear, plasmid and genomic DNA were purified using the Zymoclean Gel DNA Recovery Kit (Zymo Research), ZymoPURE Plasmid Miniprep Kit (Zymo Research) and EZ1 DSP Virus Kit (Qiagen), respectively.

For heat-shock transformation of *E. coli* DH10B or DH5α, CaCl_2_-competent cells were mixed with 10% (v/v) of ligation reactions and incubated on ice for 15 min. Cells were then heat-shocked at 42°C for 40 s, immediately chilled on ice, and recovered with 900 µL of SOC (2% tryptone, 0.5% yeast extract, 10 mM NaCl, 2.5 mM KCl, 10 mM MgCl_2_, 10 mM MgSO_4_ and 20 mM glucose; pH 7) for 1 h before plating in selective medium.

### Construction of a fluorescence-luminescence transcriptional reporter

A DNA cassette containing a *hlpA-mNeonGreen* fusion and plasmid pBIR, which integrates in the genome of *S. mitis* G22 and *S. pneumoniae* D39, were synthesized (Twist Bioscience). The reporter cassette and plasmid were amplified and blunt end ligated to generate plasmid pJB10. The firefly luciferase *luc* (from pPEP-LGZ) and plasmid pJB10 were amplified and blunt-end ligated to create plasmid pJB11 encoding a dual fluorescence-luminescence reporter. Transcriptional fusions were created by digesting both pJB11 and promoter fragments with BsaI, followed by ligation. Variants of pJB11 were linearized with BamHI and transformed into *S. mitis* G22 or *S. pneumoniae* D39. Transformants were selected in BA with chloramphenicol.

### Expression of *cabABC* and other competence genes

*S. mitis* and *S. pneumoniae* strains harboring transcriptional fusions of competence promoters were grown in C+Y_YB_ at pH 6.6 (to prevent natural competence induction) until an OD_600_ of 0.5. To evaluate natural competence dynamics, acidic precultures were diluted 1:100 in C+Y_YB_ (pH 6.6, 7.4 or 8.2) containing 340 µg/mL D-luciferin (Abcam). To assess the effect of different pheromones on competence induction, acidic precultures were diluted to OD_600_ = 0.1 in C+Y_YB_ (pH 6.6) with 340 µg/mL D-luciferin. Then, CSP or BlpC variants (up to 1 μM), were added. Cultures were transferred to white 96-well plates with clear bottoms (Nunc). The OD_595_ and luminescence was recorded every 10 min for 24 h in a BioTek Neo2 Multimode Reader with a gain of 185. Luminescence values were normalized to the OD_595_. D-luciferin was resuspended in sterile ddH_2_O to 15 mg.mL^-1^ and kept at-80 °C in single-use aliquots.

### Construction of ComE, SigX1 and SigX2 deletion mutants

The upstream and downstream flanking regions of *comE, sigX1, sigX2* were amplified from *S. mitis* G22 gDNA. Kanamycin and spectinomycin resistance genes were amplified from pKan and pPEP-LGZ (70), respectively. DNAs were ligated using Gibson Assembly (New England Biolabs) and amplified with nested primers. Transformants were selected in BA with kanamycin (for *comE* and *sigX1*) or spectinomycin (for *sigX2*).

### Biofilms and transwells

*S. mitis* and *S. pneumoniae* were grown to exponential phase and mixed at a 1:1 ratio (10^5^ CFU of each strain) in C+Y_YB_ with 1600 U.mL^-1^ catalase. Two milliliters of the mixtures were added to tissue-culture treated 12- or 24-well plates (Corning) and incubated 24 h at 34°C with 5% CO_2_. Non-adherent cells were removed by aspiration, and biofilms were thoroughly resuspended in PBS. Initial biofilms were performed in 24-well plates, while the remaining were performed in 12-well plates (since these were the ones where we could fit a transwell insert).

For transwell biofilms, 1.5 mL of C+Y_YB_ containing catalase and 10^5^ CFU of *S. pneumoniae* was added to wells of tissue-culture treated 12-well plates. Transwell inserts containing a 0.4 µm pore polycarbonate membrane (Corning) were placed in the wells, and 0.5 mL of C+Y_YB_ with catalase and 10^5^ CFU of *S. mitis* were added to the upper insert. Biofilms were recovered from both compartments.

In dual-species biofilms, *S. pneumoniae* and *S. mitis* cells were enumerated by plating on BA with chloramphenicol or kanamycin, respectively. In the transwell setup, where *S. pneumoniae* and *S. mitis* were physically separated, biofilm suspensions from both compartments were also plated on selective media to confirm that neither strain had crossed the membrane.

### Construction of an *S. mitis cab* overexpressor mutant

The plasmid backbone was amplified from pDL278_P23-DsRed-Express2 (Addgene plasmid #121505), and *cabABC* was amplified from *S. mitis* G22 gDNA. The two fragments were blunt-end ligated, generating plasmid pJB09-*cabABC*, in which *cabABC* is placed under the control of the strong constitutive P23 promoter. *S. mitis* transformants were selected in BA with spectinomycin.

### Inhibitory activity of cell-free supernatants and heat-killed cells

*S. mitis* was grown to early stationary phase and centrifuged at 10,000 g for 30 min at 4°C. Cell-free supernatants (CFS) were obtained by filtration (0.22 µm). CFS were lyophilized by vacuum evaporation using a SpeedVac concentrator (LabConco) and resuspended in ddH_2_O to a concentration of 10x before testing. C+Y_YB_ medium was concentrated in parallel and used as a negative control for inhibitory activity. Cell pellets were washed once in PBS and heat-killed for 15 min at 95 °C. Complete lysis of pellet extracts was confirmed by plating in BA.

To test for inhibitory activity of CFS and heat-killed cells, *S. pneumoniae* D39 was grown to an OD_600_ of 0.5, diluted 1:100 in fresh medium, and 180 µL of the dilution was added to clear 96-well plates. Twenty microliters of treatments or water was added to each well and growth was monitored by measuring the OD_595_ every 30 min for 24 h using a Tecan Infinite 200 Pro.

### Sensitivity of bacteria to CabAB

Exponential-phase cultures of bacterial species (18 *Streptococcus* spp., four additional non-streptococcal Gram-positive species, and four Gram-negative species) were diluted to OD_600_ 0.1 and 195 µL was added to wells of clear 96-well plates. CabAB (0.975 µM) or DMSO (0.00975%) was added to wells and growth was recorded by measuring the OD_595_ every 10 min for 24 h using the Tecan Infinite 200. Growth arrest was quantified by comparing the area under the curve (AUC) between CabAB- and DMSO-treated cultures. All further assays using CabAB and DMSO treatments were performed in the same conditions, unless specified otherwise.

### Construction of autolysis-deficient *S. pneumoniae* mutants

The upstream and downstream regions of *lytA* and *lytC* were amplified from *S. pneumoniae* D39 gDNA. The PhunSweet cassette containing *pheS-sacB-*aph*A-3* which confers susceptibility to chlorinated phenylalanine and sucrose, and resistance to kanamycin, was amplified from *S. pneumoniae* D39 Δ*cps*::*PhunSweet* gDNA (71). DNAs were assembled using Gibson Assembly and amplified with nested primers. *S. pneumoniae* D39 transformants were selected on BA with kanamycin.

To remove the antibiotic resistance cassette, the flanking regions of *lytA* and *lytC* were amplified, assembled with Gibson Assembly, and amplified with nested primers. Transformants were recovered by counterselection on BA with 15 mM chlorinated phenylalanine and 10% sucrose.

### Scanning electron microscopy (SEM)

*S. pneumoniae* D39 cultures (30 mL) were treated with DMSO or CabAB and incubated for 15 min at 37°C. Cells were harvested by centrifugation (all centrifugations were carried out at 800 g for 10 min at room temperature) and washed three times with PBS. Fixation was performed in 500 μL of 2.5% glutaraldehyde and 1% paraformaldehyde in 0.1 M sodium cacodylate buffer for 30 min at room temperature. Cells were washed three times with 0.1 M sodium cacodylate buffer and dehydrated in increasingly higher ethanol concentrations (50%, 70%, 90%, 100%), by incubating 10 min in 500 μL of each ethanol concentration. Ethanol was replaced with 500 μL tert-butyl alcohol and samples were incubated at 30°C for 1 h. Then, tert-butyl was removed and replaced with 500 μL of fresh tert-butyl alcohol. Samples were kept at-20°C overnight. Samples were then lyophilized under vacuum, gold-sputtered using a Cressington 108 sputter coater, and imaged with a Hitachi SU-8010 scanning electron microscope at 1.5kV.

### Transmission electron microscopy (TEM)

*S. pneumoniae* D39 cultures (30 mL) were treated with DMSO or CabAB, incubated for 15 min at 37°C, and recovered by centrifugation at 800 g. Cells were fixed with 2% formaldehyde, 2.5% glutaraldehyde, 0.075% ruthenium red and 1.55 % lysine acetate in distilled water for 20 min at 4°C. Cells were washed twice in 0.075% ruthenium red, resuspended in 2% formaldehyde, 2.5% glutaraldehyde and 0.075% ruthenium red, and incubated overnight at 4°C. Cells were washed three times with 0.075% ruthenium red and further processed at Instituto Gulbenkian de Ciência. Cells were embedded in 2% low-melting point agarose and fixed with 1% osmium tetroxide and 0.06% ruthenium red for 1 h at room temperature. Cells were washed once with 0.1 M sodium cacodylate buffer and twice with distilled water. Dehydration was performed on ice using increasing concentrations of ethanol (10%, 30%, 50%; 10 min each). Samples were stained overnight at 4°C with 2% uranyl acetate in 70% ethanol, followed by further dehydration with 90% ethanol and three times with 100% ethanol. Infiltration was carried out with increasing concentrations of epoxy resin in ethanol (3:1, 1:1, 1:3), and 100% resin overnight, followed by two exchanges of 100% resin. Ultrathin sections were cut and imaged on a Tecnai G2 Spirit BioTWIN Transmission Electron Microscope (FEI) equipped with an Olympus-SIS Veleta CCD camera.

Large intracellular aggregates were observed in some CabAB-treated cells. To quantify these structures, micrographs of CabAB- and DMSO-treated cells were blindly evaluated by two researchers. Each researcher was first shown representative cells with and without aggregates, and then independently counted the number of cells containing these structures. The final percentage of cells with aggregates was calculated as the average between both researcher counts.

### Permeabilization and depolarization of *S. pneumoniae* membranes

*S. pneumoniae* D39 cultures were treated with DMSO or CabAB. One-milliliter aliquots were taken at 0, 2, 5, 15, 30, 60, and 120 min post treatment and immediately chilled on ice. Cells were washed twice with PBS and kept on ice. Cells were stained with 6 µM propidium iodide (Invitrogen) and 3 µM DiBAC_4_(3) (Enzo Life Sciences) for 5 min at room temperature, protected from light. Cells were centrifuged, resuspended in PBS, and added to black 96-well plates with clear bottoms (Nunc). Fluorescence intensity was measured using a BioTek Neo2 Multimode Reader under the following settings: shake for 5 s; propidium iodide – top reading, 535 ± 12 nm excitation, 617 ± 12 nm emission and 120 gain; DiBAC_4_(3) – top reading, 492 ± 9 nm excitation, 518 ± 9 emission, 120 gain). Fluorescent values were normalized to the OD_595_.

### Efflux of ATP

Cultures of *S. pneumoniae* were treated with DMSO or CabAB. Aliquots were taken at 0, 15 and 120 min post-treatment and kept on ice. Cells were centrifuged, and supernatants were filtered (0.22 µm) and maintained on ice. Cells were permeabilized in DMSO for 10 min at room temperature. Both cells and supernatants were diluted 1:10 in ddH_2_O and kept on ice.

Extracellular and intracellular ATP was measured with the ATP Luminescent Cell Viability Assay (Millipore) according to instructions. Luminescence was measured in a BioTek Neo2 Multimode Reader using a gain of 150 and integration time of 10 s. An ATP standard curve confirmed linear detection within the range of 0.2 μM to 2 mM ATP. Total ATP was calculated by summing luminescence values from both the cell and supernatant fractions under each condition.

### Construction of *S. mitis* harboring HA-tagged bacteriocins

DNAs containing HA (YPYDVPDYA)-tagged *cabA* and *cabB* were synthesized (Twist Bioscience). The regions upstream (including the native promoter) and downstream (including a chloramphenicol resistance cassette) of the *cab* operon were amplified from pJB11-*cabABC*. DNAs were assembled by Gibson Assembly and amplified with nested primers. The resulting constructs were transformed into the *S. mitis* Δ*cabABC* background, generating two mutant strains: *cabA-HA cabBC* and *cabA cab-HA cabC.* Transformants were selected in BA with chloramphenicol.

### Confocal microscopy and immunofluorescence

To quantify the percentage of cells that become membrane-permeable, *S. pneumoniae* was treated with DMSO or CabAB. Cells were recovered by centrifugation (9600 g, 4°C, 3 min) and washed three times with PBS. Cells were stained with 30 µM propidium iodide and 5 µM SYTO 9 for 15 min at room temperature, protected from light.

To determine whether large intracellular structures observed in CabAB-treated *S. pneumoniae* cells corresponded to membranes, treated cells were washed three times with PBS and stained with 20 µg.mL^-1^ FM4-64 and 15 µg.mL^-1^ DAPI for 5 min at room temperature, protected from light.

For immunofluorescence of live competent bacteria, *S. mitis* harboring HA-tagged bacteriocins were grown in acidic C+Y_YB_ and diluted to OD_600_ = 0.1 in pre-warmed C+Y_YB_. CSP_G22_ was added at 1μM, and cells were incubated for 70 min at 37°C and then chilled on ice. To retain the competent status of cells, all subsequent steps were performed on ice using ice-cold solutions. Cells were centrifuged (9600 g, 4°C, 3 min) and washed twice in PBS. Blocking was performed in PBS with 5% bovine serum albumin (BSA) for 1 h. Cells were pelleted and resuspended in 100 µL of primary rabbit monoclonal anti-HA antibody (clone C29F4; Cell Signaling Technology) diluted 1:100 in PBS with 1% BSA, and incubated for 1 h. Cells were washed twice in PBS with 0.1% Tween 20, resuspended in 100 µL of Alexa Fluor 488-conjugated goat anti-rabbit secondary antibody (Invitrogen) diluted 1:500 in PBS with 1% BSA, and incubated 1 h protected from light. Cells were washed twice with PBS containing 0.1% Tween 20 and once with PBS. Cells were then stained with 20 µg.mL^-1^ FM4-64 and 15 µg.mL^-1^ DAPI for 5 at room temperature, covered from light.

Immediately before imaging, cells were mounted on thin 1.7% agarose pads. Fluorescence imaging was performed using a Zeiss LSM 880 point scanning confocal microscope equipped with an Airyscan detector. Samples were imaged with a Plan-Apochromat 63x/1.4 Oil DIC M27 objective (Zeiss), using laser lines 405 nm (DAPI), 488 nm (SYTO 9 and Alexa Fluor 488), and 561 nm (FM4-64 and propidium iodide). Imaging was performed at laser intensities of 2-3% and a gain of 700 for DAPI, 700 for SYTO 9, 1000 for Alexa Fluor 488, 900 for FM4-64, and 700 for propidium iodide. The Zeiss Zen v3.0 was used to control the microscope and for Airyscan processing. Image analysis was conducted using ImageJ/FIJI v2.14 (72).

### Construction of a transformation-deficient *S. pneumoniae* mutant

The upstream and downstream regions of *comGC* were amplified from *S. pneumoniae* D39 gDNA, and *tetM* was amplified from *S. mitis* F22 gDNA. DNAs were ligated by Gibson Assembly and amplified using nested primers. *S. pneumoniae* transformants were selected in BA with tetracycline.

### Horizontal gene transfer in biofilms

Dual-species biofilms of *S. pneumoniae* D39-GFP Δ*comGC* (chloramphenicol and tetracycline resistant; Chl^R^ Tet^R^) and *S. mitis* (kanamycin resistant; Kan^R^) were grown as described above, except that C+Y_YB_ was adjusted to pH 6.6 to prevent natural competence induction. After 24h, non-adherent cells were removed by aspiration, and 2 mL of fresh C+Y_YB_ (pH 7.4) containing catalase and 1 µM of CSP_G22_ were added to biofilms. After 6 h of incubation at 34°C with 5% CO_2_, biofilm cells were thoroughly resuspended in PBS and recovered by centrifugation.

Cells were plated on the following selective media: (i) BA with kanamycin to quantify total *S. mitis* (Kan^R^); (ii) BA with kanamycin and chloramphenicol to detect *S. mitis* transformants that have acquired chloramphenicol resistance (Kan^R^ Chl^R^), either with or without tetracycline resistance; (iii) BA with kanamycin and tetracycline to detect *S. mitis* transformants that acquired tetracycline resistance (Kan^R^ Tet^R^), either with or without chloramphenicol resistance; (iv) BA with kanamycin, chloramphenicol and tetracycline to identify *S. mitis* transformants that acquired both resistances (Kan^R^ Chl^R^ Tet^R^); (v) and BA with chloramphenicol and tetracycline (without kanamycin) to detect *S. pneumoniae* loads.

Transformants were counted after 48 h of incubation. *S. mitis* transformation frequencies were calculated as the ratio between the number of transformants (Kan^R^ colonies also resistant to chloramphenicol and/or tetracycline) and the total number of Kan^R^ colonies. *S. mitis* G22 Δ*comE* was used as a non-transformable control, in which acquisition of chloramphenicol and tetracycline resistance should not occur via transformation.

## Statistical analyses

The two-tailed unpaired *Mann-Whitney U* test with Benjamini and Hochberg correction for FDR was used to compare bacterial loads. Student’s *t*-test with Benjamini and Hochberg correction for FDR was used to compare gene expression. The ratio paired *t*-test was used for the statistical analysis of bacterial proportions in multi-species biofilms. All statistical analysis were computed using GraphPad Prism 9.0 and considered significant at *P*-value ≤0.05.

## Data availability

All data is available from the corresponding author upon reasonable request.

## Acknowledgements

We are grateful to Jan-Willem Veening (University of Lausanne) and Morten Kjos (Norwegian University of Life Sciences) for providing plasmids pPEP-LGZ and Jason Rosch (St. Jude Children’s Research Hospital) for providing *S. pneumoniae* strain harboring a PhunSweet cassette. We also acknowledge Luisa N. Hiller (Carnegie Mellon University) and Adriano O. Henriques (ITQB NOVA) for fruitful discussions.

## Author Contributions

R. Sá-Leão and J. Borralho contributed to the concept and design of the study. R. Sá-Leão contributed with reagents and materials. Data acquisition was performed by J. Borralho, J. Lança, J. Bryton and W. Antunes. Data was interpreted by J. Borralho, J. Lança, J. Bryton, W. Antunes and R. Sá-Leão. The manuscript was drafted by J. Borralho and R. Sá-Leão. All authors critically revised and approved the final version of the manuscript.

## Supplemental material

**Fig. S1.**
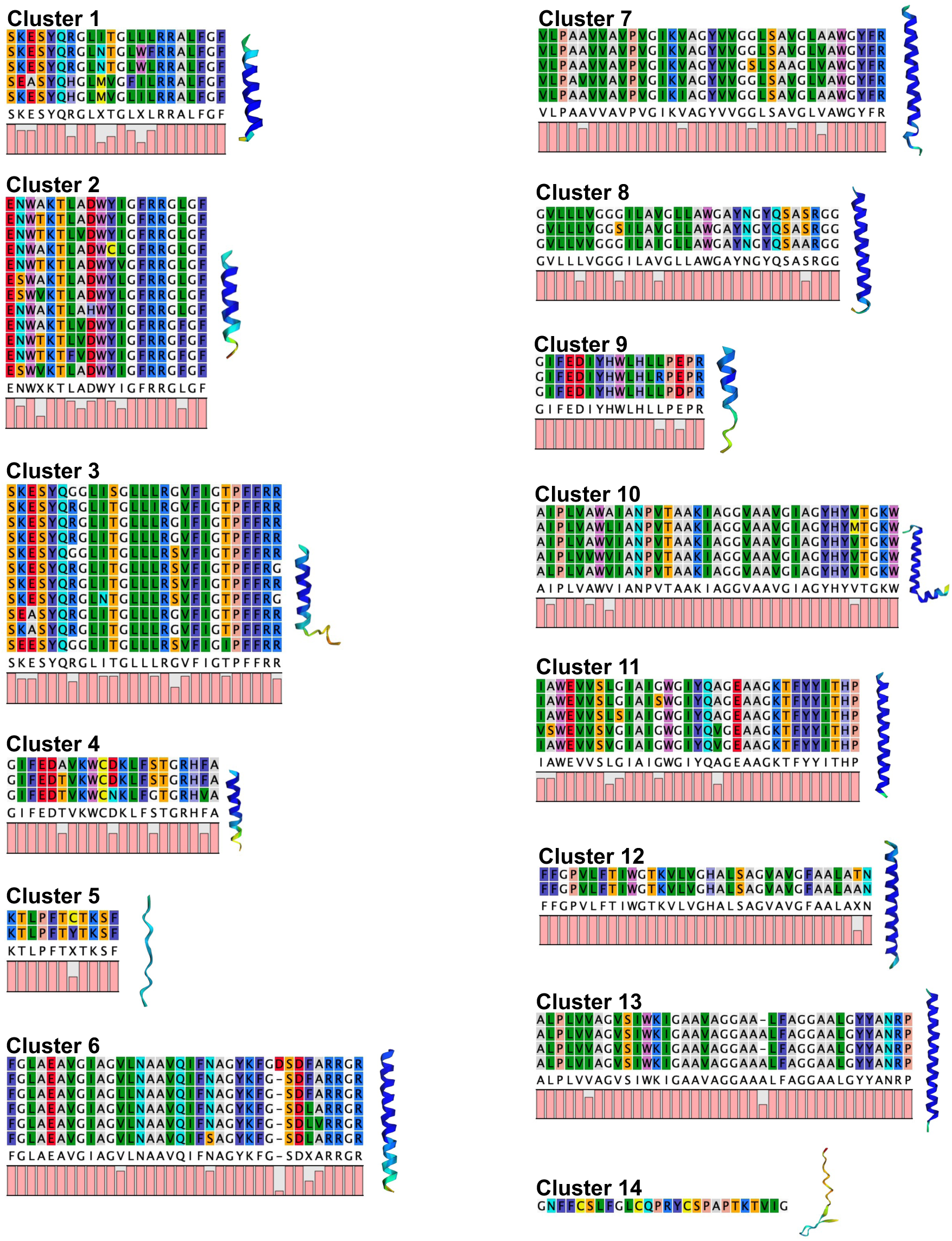
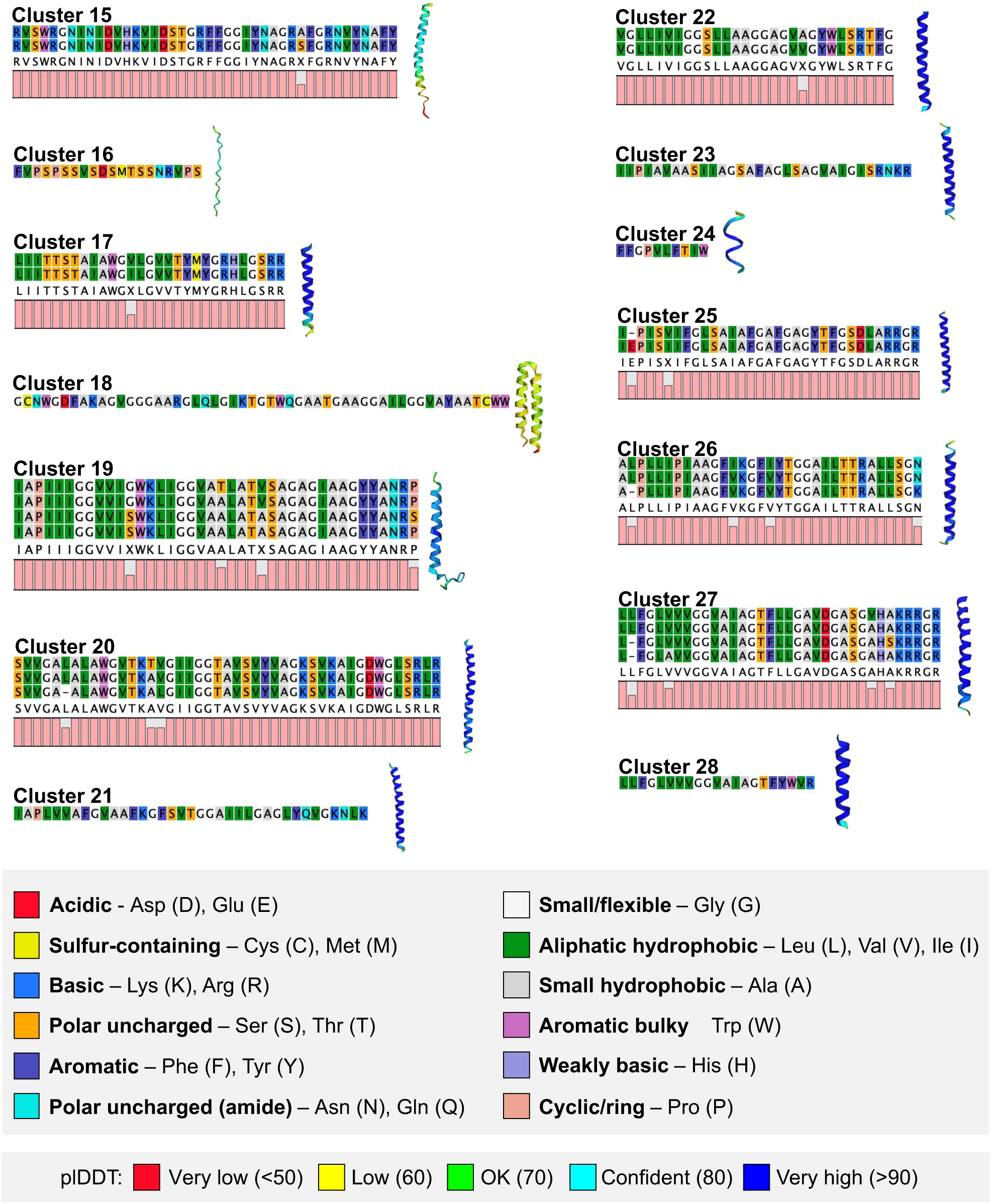
Alignment-based clustering of Cab variants. The 95 Cab variants found *S. mitis* populations were clustered based on pairwise sequence alignments (see Materials and Methods). Alignments for Cab sequences within each cluster is shown, with associated Alphafold2 (ColabFold v1.5.5) structural model of the most prevalent variant. Amino acids are colored according to biochemical traits (RasMol coloring). Structures are colored according to plDDT (predicted local distance difference test) confidence.

**Fig. S2.**
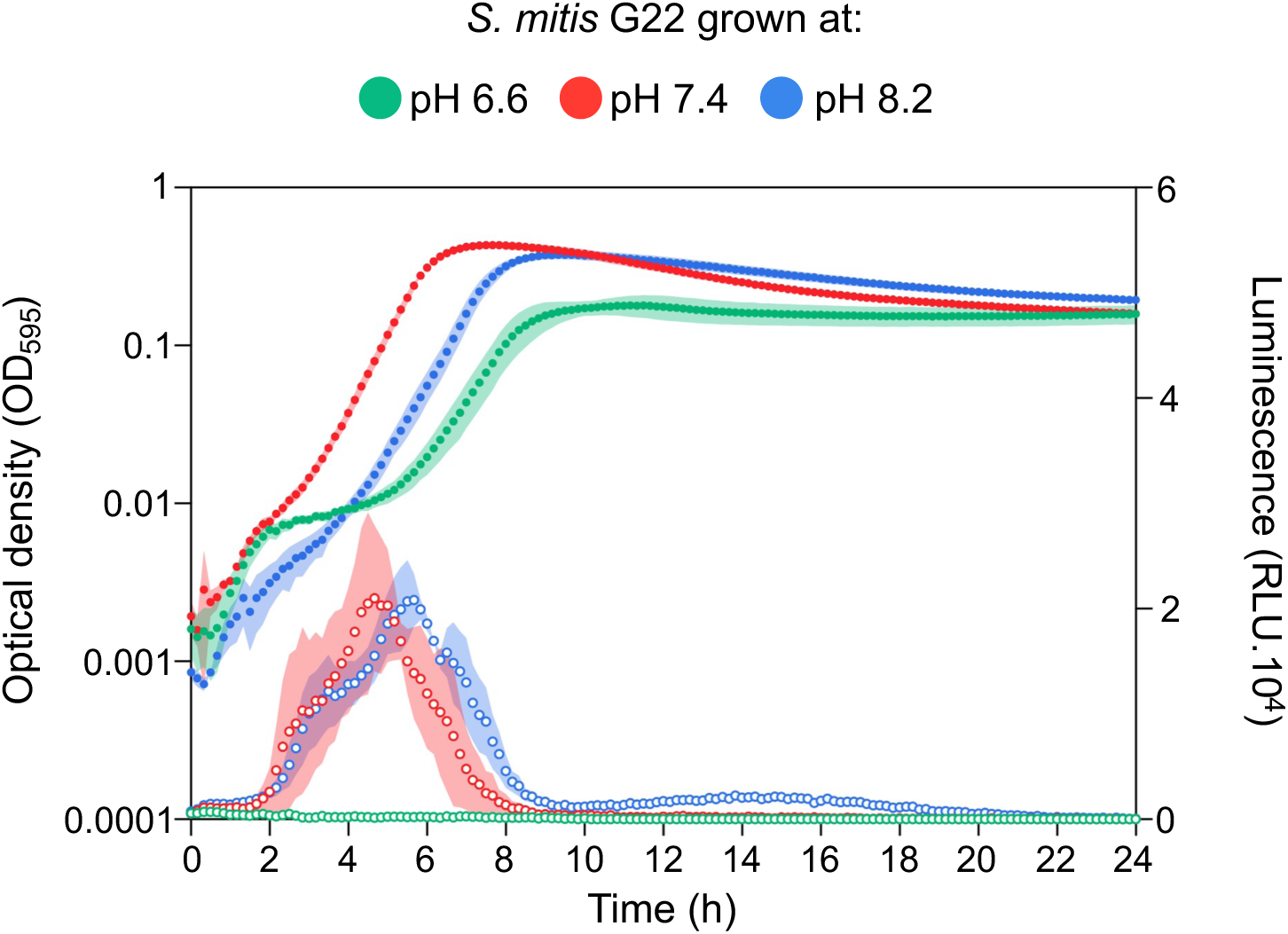
*S. mitis* competence is inhibited in medium with acidic pH. *S. mitis* G22 harboring an early-competence reporter (P*comCDE*-*luc*) was pre-grown in medium at pH 6.6, before being diluted 1:100 in fresh medium at pH 6.6, 7.4, or 8.2. Optical density (closed circles) and luminescence (open circles) values were measured every 10 min for 24 h. Growth curves were repeated three times independently, and one representative experiment with two technical replicates per condition is shown.

**Fig. S3.**
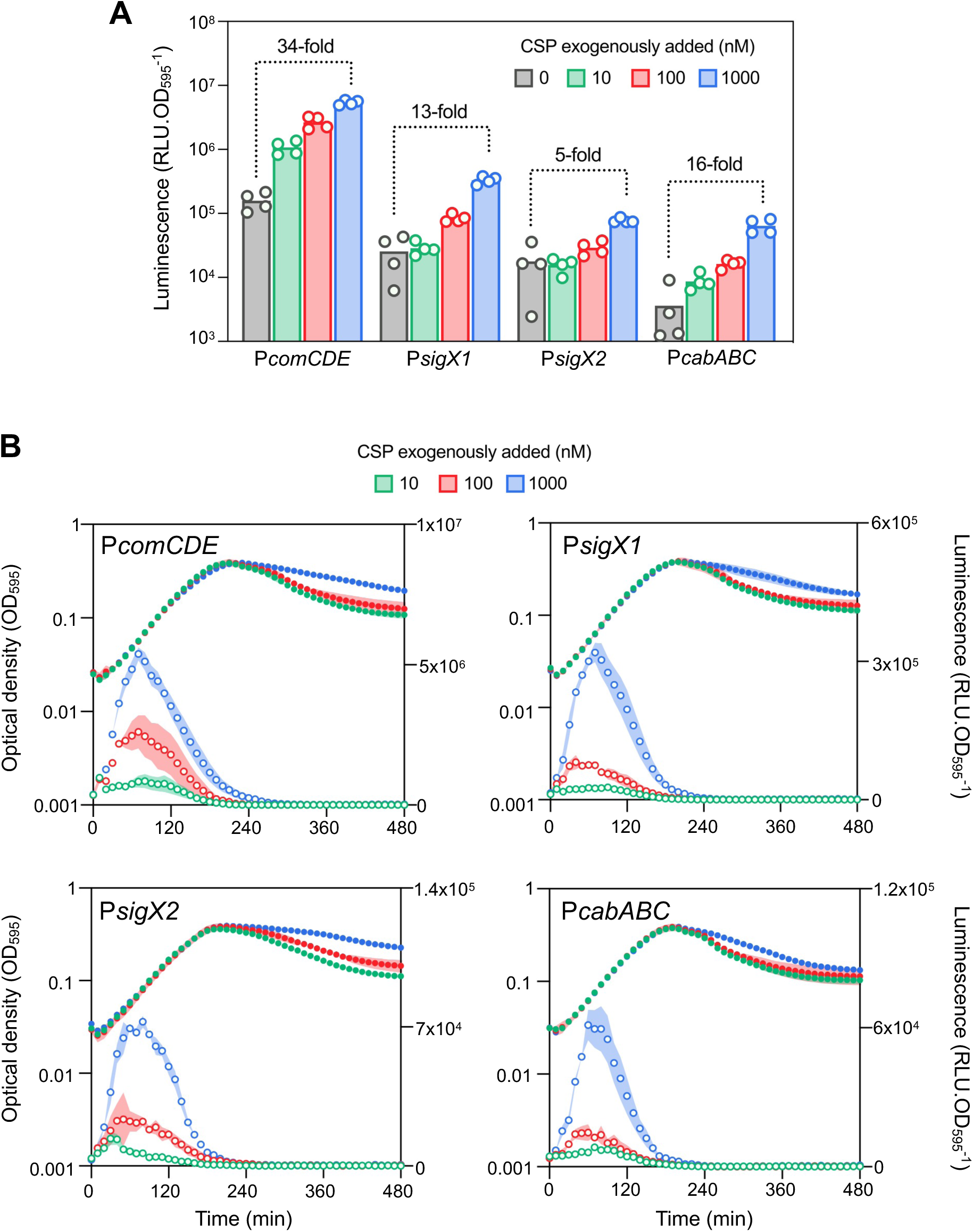
Expression of *cabABC* follows the same dynamics of other competence genes. (A) Expression of *comCDE*, *sigX1*, *sigX2* and *cabABC* evaluated by luminescence produced by luminescent reporter *S. mitis*. Cells were pre-grown in acidic pH and diluted in fresh medium containing synthetic CSP_G22_. Values represent the highest expression value (RLU.OD_595_^-1^) obtained during the 3 h post CSP_G22_ treatment. (B) Dynamics of *comCDE*, *sigX1*, *sigX2* and *cabABC* expression in *S. mitis*. Optical density – closed circles. Luminescence – open circles. Growth curves were repeated four times independently, and one representative experiment with two technical replicates per condition is shown. For clarity only the first 8 h of 24 h are shown, since there is no visible expression of the four reporters after this period.

**Fig. S4.**
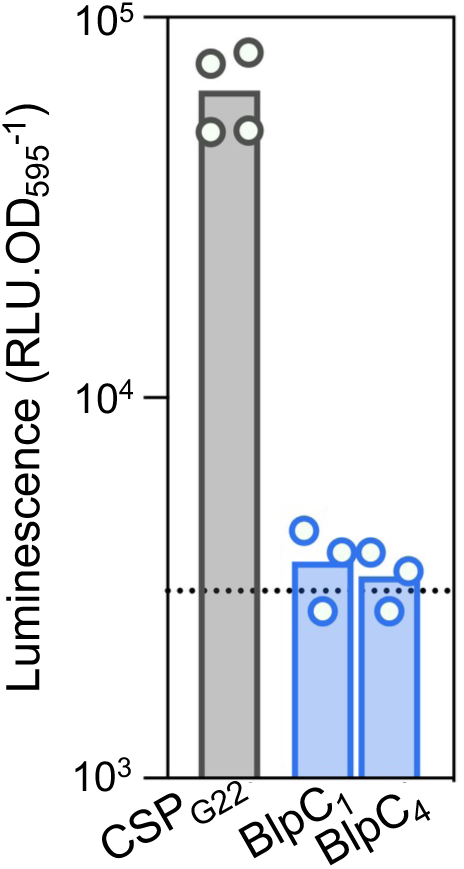
Expression of *cabABC* is not induced by bacteriocin-like peptide pheromones (BlpC). *S. mitis* G22 harboring a P*cabABC*-*luc* reporter was pre-grown in acidic pH and diluted in fresh medium containing 1 µM of synthetic CSP_G22_ (grey), 1 µM of BlpC_1_ or BlpC_4_ pheromones (blue) or left untreated (dashed line). BlpC_1_ or BlpC_4_ are encoded in two *blp*-type bacteriocin clusters elsewhere in the genome of G22. Experiment were repeated four times independently.

**Fig. S5.**
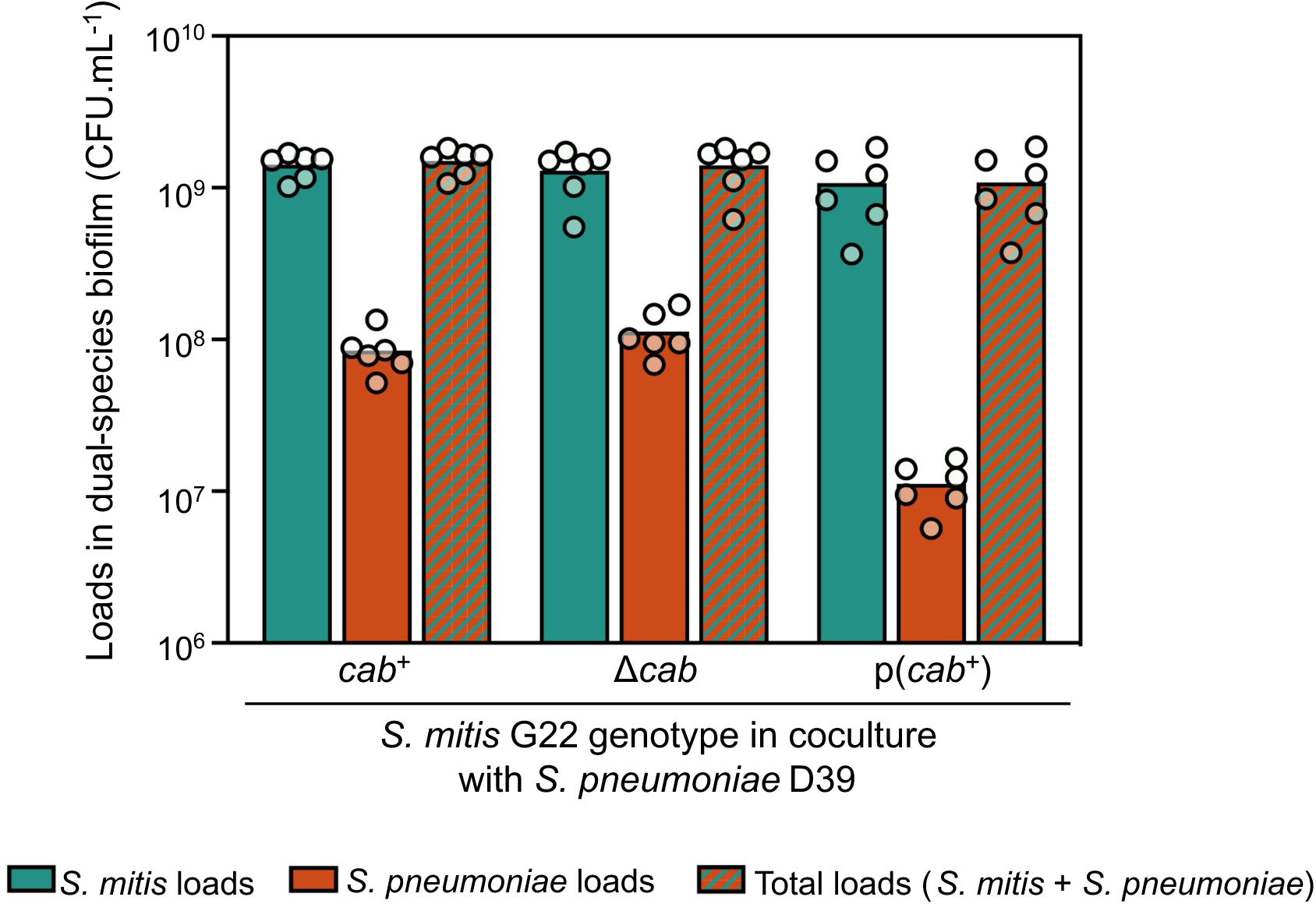
Expression of *cab* by *S. mitis* reduces *S. pneumoniae* loads in dual-species biofilms. Individual and total bacterial loads of biofilm proportions presented in Figure 4A of the main text. *S. pneumoniae* and *S. mitis* of different *cab* genotypes were grown in 24 h biofilms in competence-permissive conditions. *S. mitis* and *S. pneumoniae* cell loads were enumerated by plating in BA with kanamycin or chloramphenicol, respectively. The experiment was repeated three times, independently.

**Fig. S6.**
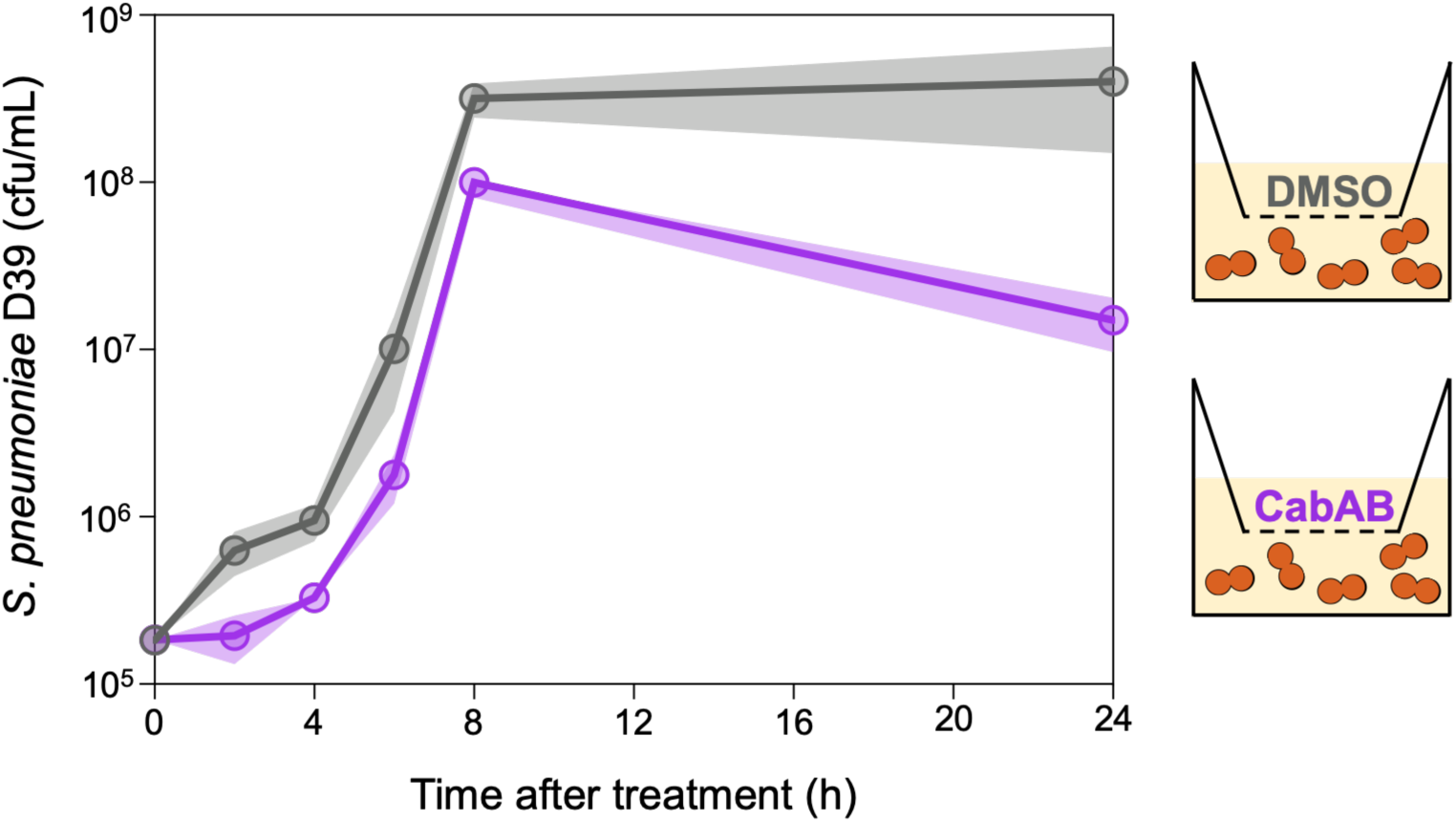
Synthetic CabAB peptides can cross the transwell membrane. *S. pneumoniae* D39 cells were diluted to ∼10^5^ cfu/mL in C+Y_YB_ and 1.5 mL was added to 12-well plates. An insert containing a 0.4 µm pore polycarbonate membrane was placed on top and 0.5 mL of C+Y_YB_ with 0.975 µM CabAB or 0.00975% DMSO was added. Aliquots were taken from the bottom compartment at indicated timepoints and plated do assess *S. pneumoniae* loads. The experiments was repeated three times, independently.

**Fig. S7.**
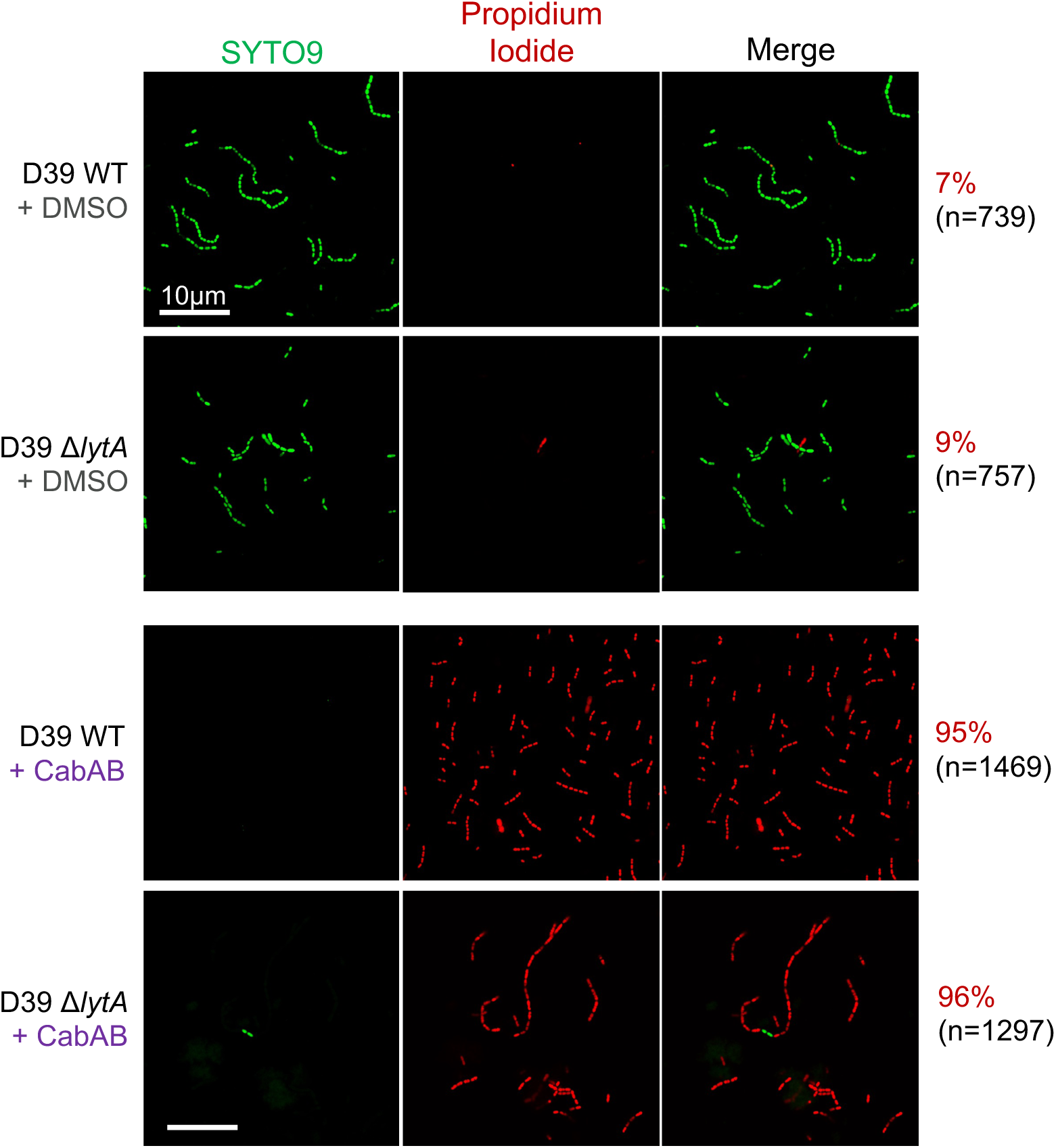
Permeabilization of *S. pneumoniae* membranes does not depend on autolysin LytA. Confocal imaging of wild-type and Δ*lytA S. pneumoniae* treated with 0.00975% DMSO or 0.975 µM CabAB peptides. Membrane permeabilization is shown in red, by entry of propidium iodide. The total number and percentage of permeable cells is shown next to panels. Microscopy images are representative of three independent experiments.

**Fig. S8.**
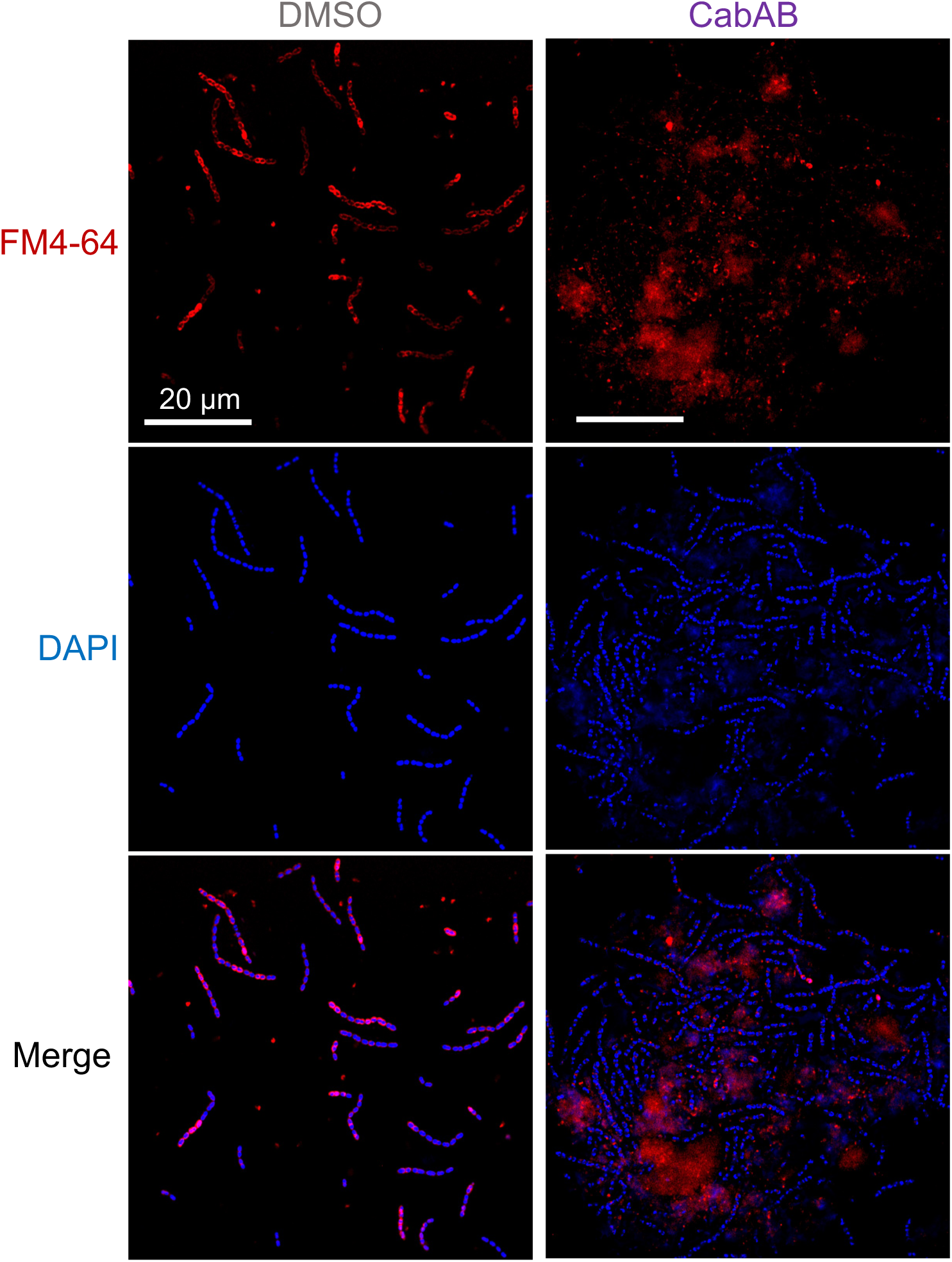
CabAB treatment induces formation of large clumps of *S. pneumoniae* cells encased in a matrix of DNA and membrane material. *S. pneumoniae* D39 was treated with 0.00975% DMSO or 0.975 µM CabAB and stained with FM4-64 (membrane dye) and DAPI (nucleic acid dye), before being imaged in a confocal microscope. Experiment was repeated three times, independently.

**Fig. S9.**
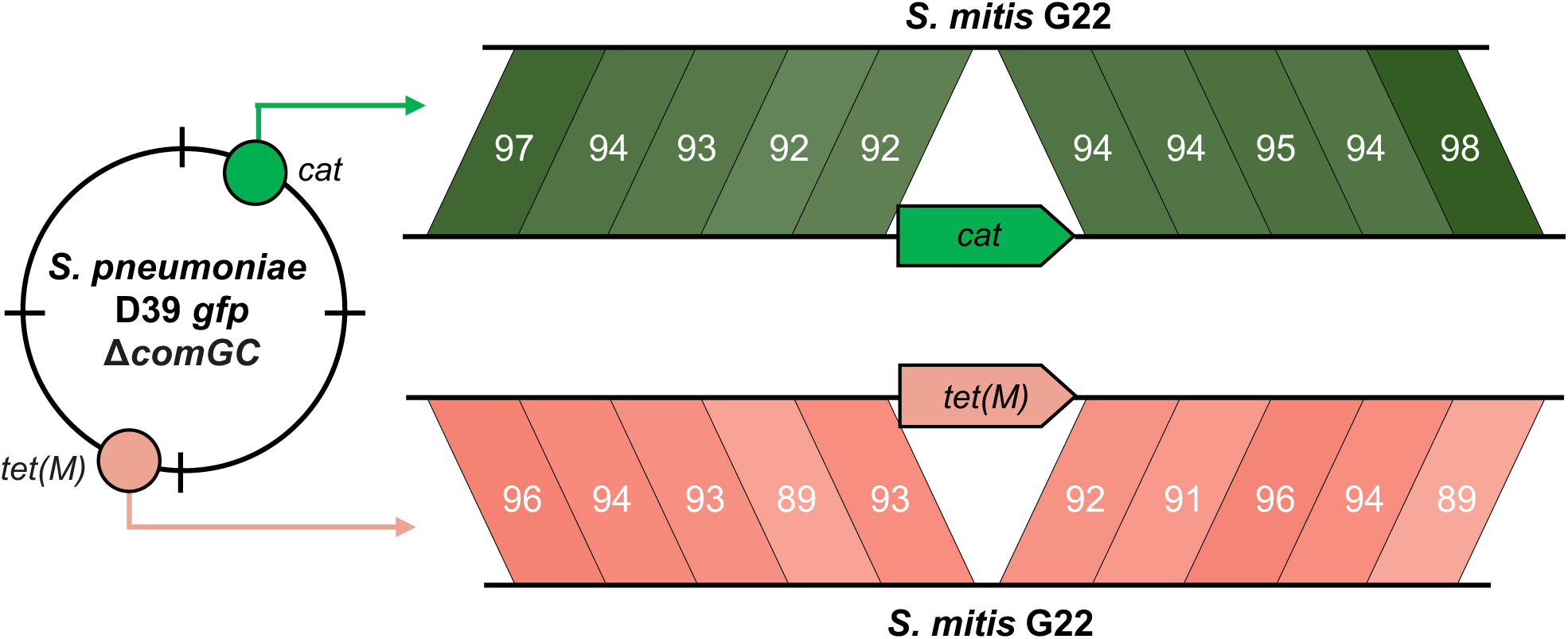
Nucleotide sequence homology between *S. pneumoniae* D39 and *S. mitis* G22 flanking the sites of antibiotic resistance gene integration. Upstream and downstream regions of *cat* and *tet(M)* in the *S. pneumoniae* DNA donor were divided in five 1,000 bp blocks each and compared to the *S. mitis* G22 genome using BLASTn. Query coverage was consistently 100%, and the percentage of sequence identity is shown for each 1,000 bp block.

**Fig. S10.**
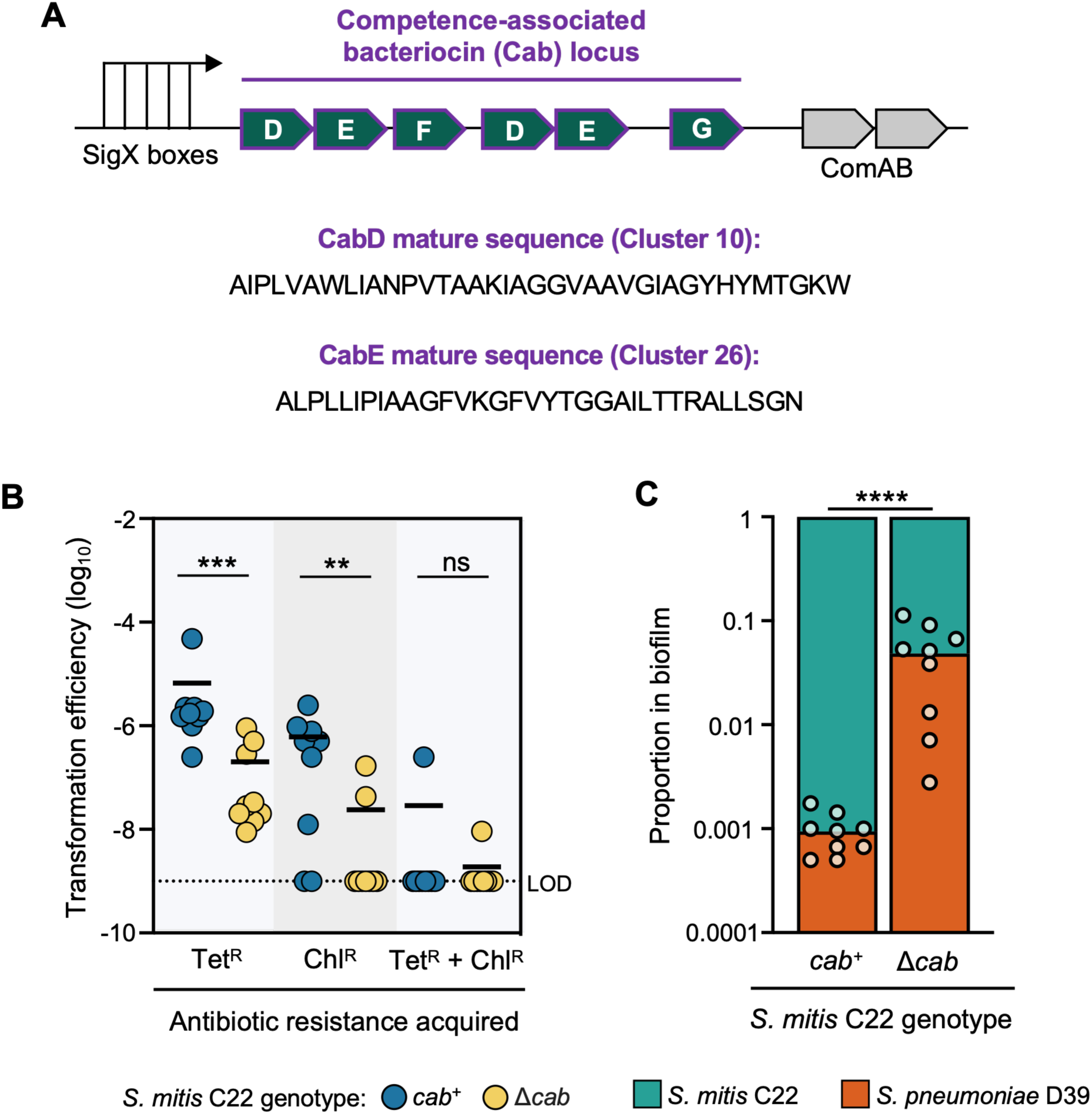
The Cab bacteriocins of *S. mitis* C22 enhance HGT and mediate inhibition of *S. pneumoniae* in dual-species biofilms. (A) Genomic arrangement of the *S. mitis* C22 *cab* locus, which contains two copies of the bacteriocin *cabD*, two copies of the bacteriocin *cabE*, and the putative immunity proteins *cabF and cabG.* (B) Acquisition of Tet^R^ and Chl^R^ by *S. mitis* C22 when in 12-well biofilms with *S. pneumoniae* DNA donor (D39 *gfp* Δ*comGC*). (C) Proportion of *S. mitis* C22 and *S. pneumoniae* D39 when grown in dual-species biofilm. Experiments were repeated three times, independently. Results in panel C represent the proportion of each species in the dual-species biofilms of panel B. Transformation efficiencies were compared using unpaired t-tests. LOD limit of detection. ** *P* < 0.01. *** *P* < 0.001. **** *P* < 0.0001.

**Fig. S11.**
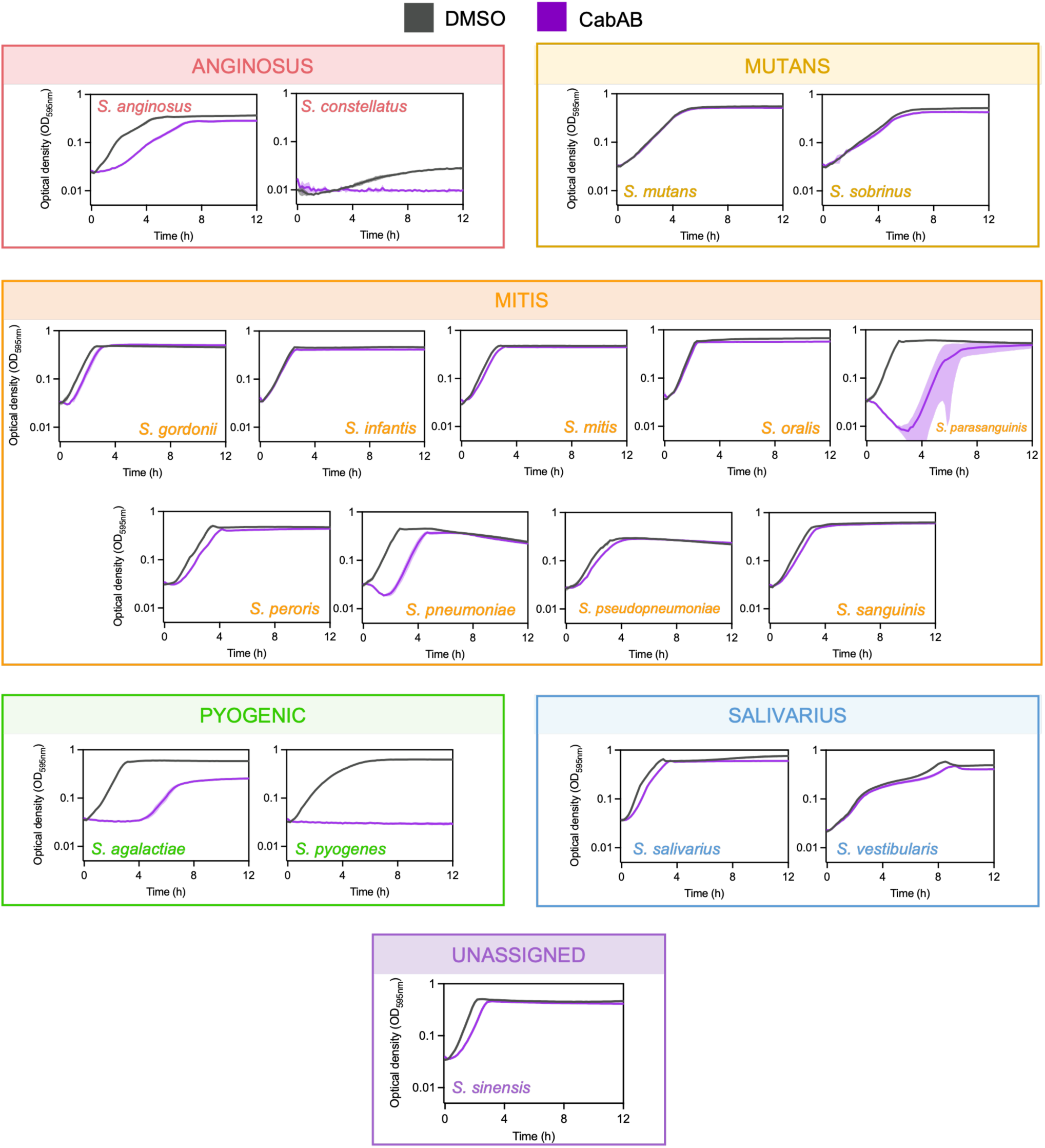
Individual growth curves of 18 streptococcal species and their sensitivity to CabAB. Exponential cultures of streptococcal species were diluted to OD_600_ 0.1 in appropriate medium and treated with either DMSO or CabAB. Optical density was measured every 10 min for 24 h. For clarity, the first 12 h are shown. Species are colored according to streptococcal group. The experiment was repeated three times, independently. Summary data of the growth curves can be found in the main manuscript.

**Fig. S12.**
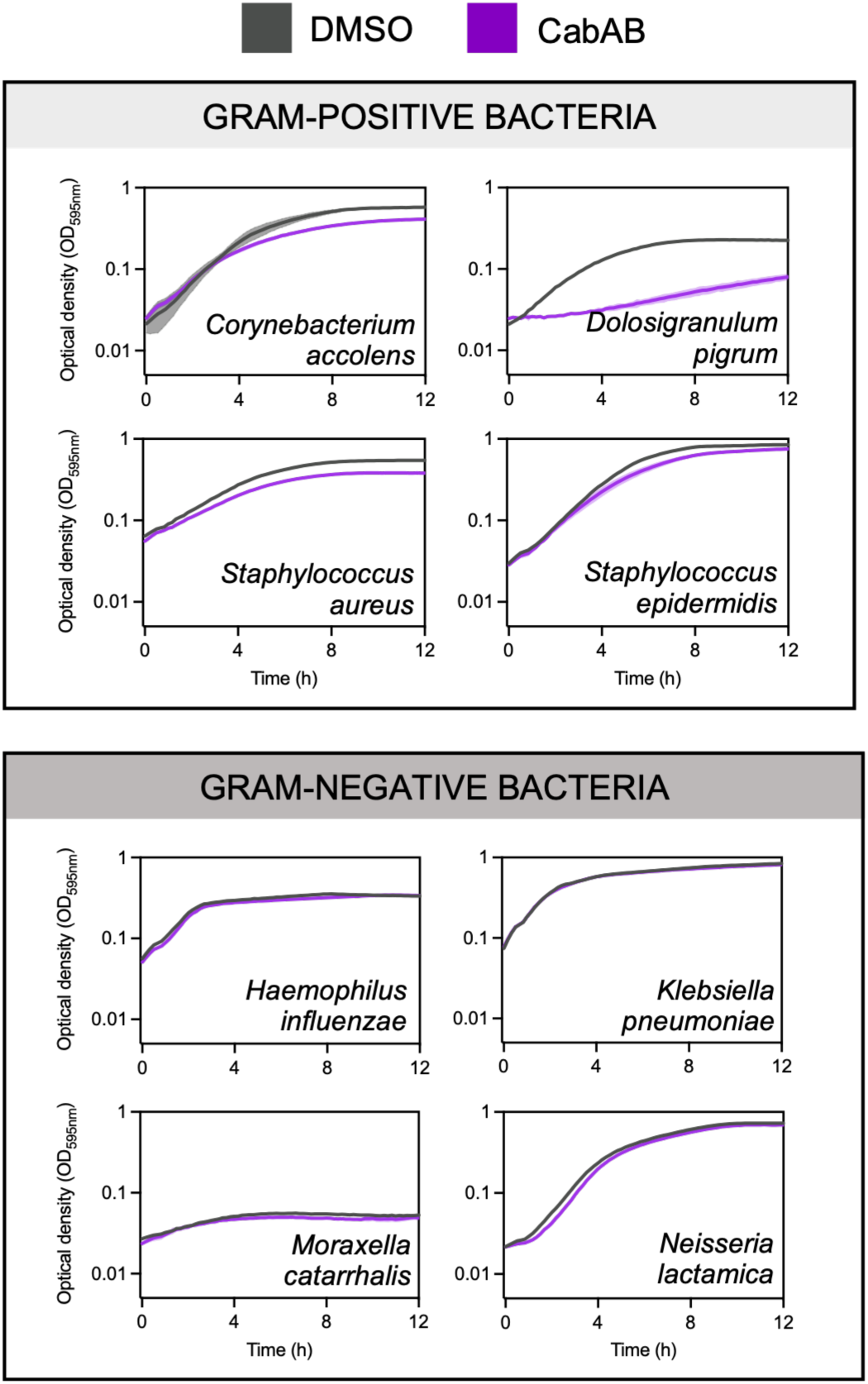
Individual growth curves of 4 non-streptococcal Gram-positive and 4 Gram-negative bacteria and their sensitivity to CabAB. Exponential cultures of species were diluted to OD_600_ in appropriate medium and treated with either DMSO or CabAB. Optical density was measured every 10 min for 24 h. For clarity, the first 12 h are shown. Species are colored according to Gram staining. The experiment was repeated three times, independently. Summary data of the growth curves can be found in the main manuscript.

**Fig. S13.**
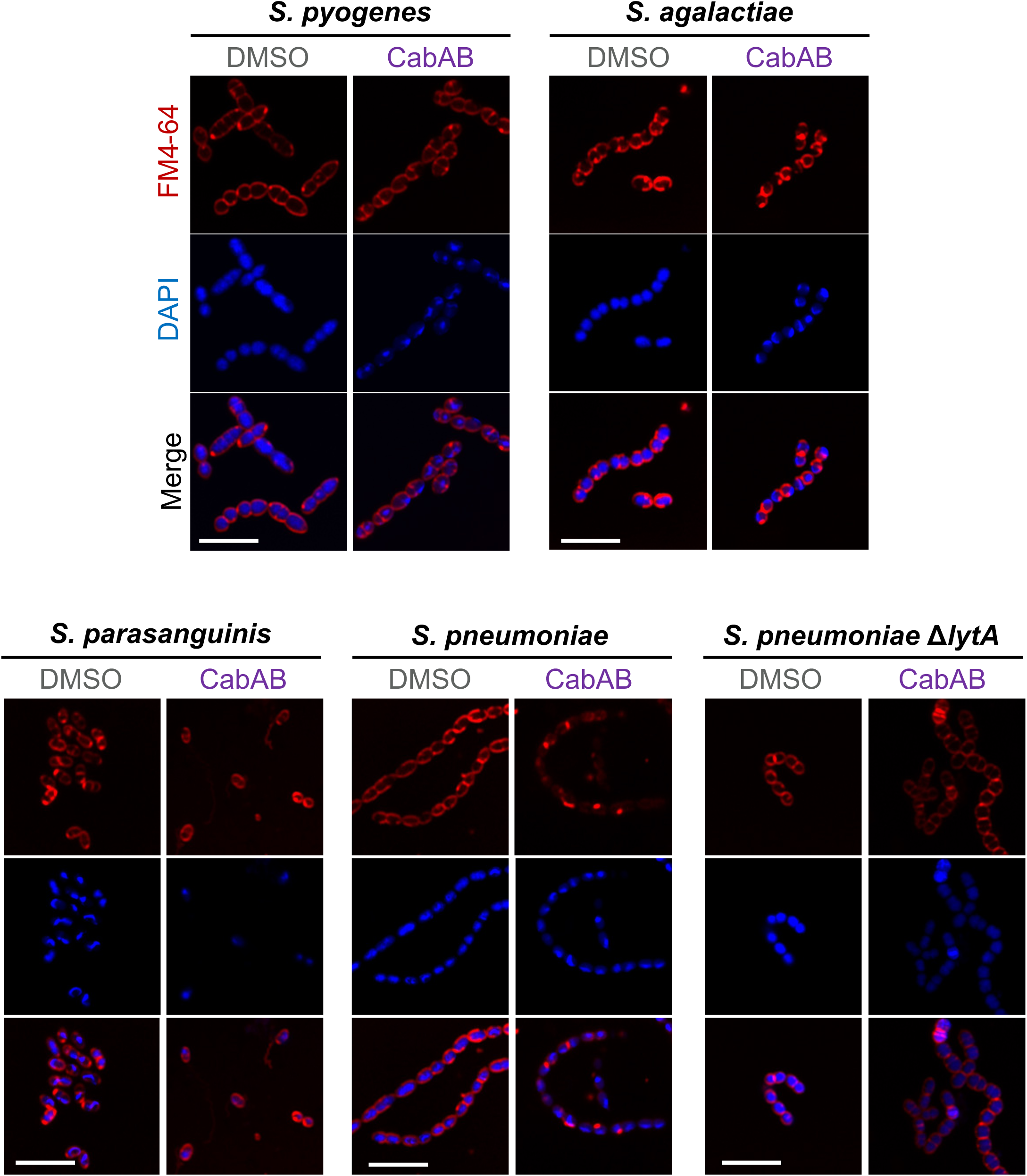
CabAB treatment induces formation of membrane aggregates exclusively in *S. pneumoniae*. Cells were treated with 0.975 µM CabAB or 0.00975% DMSO for 15 min and stained with FM4-64 (membrane dye) and DAPI (nucleic acid dye), before being imaged in a confocal microscope. The experiment was repeated three times, independently. Scale – 5 µm.

---

**Table S1.**
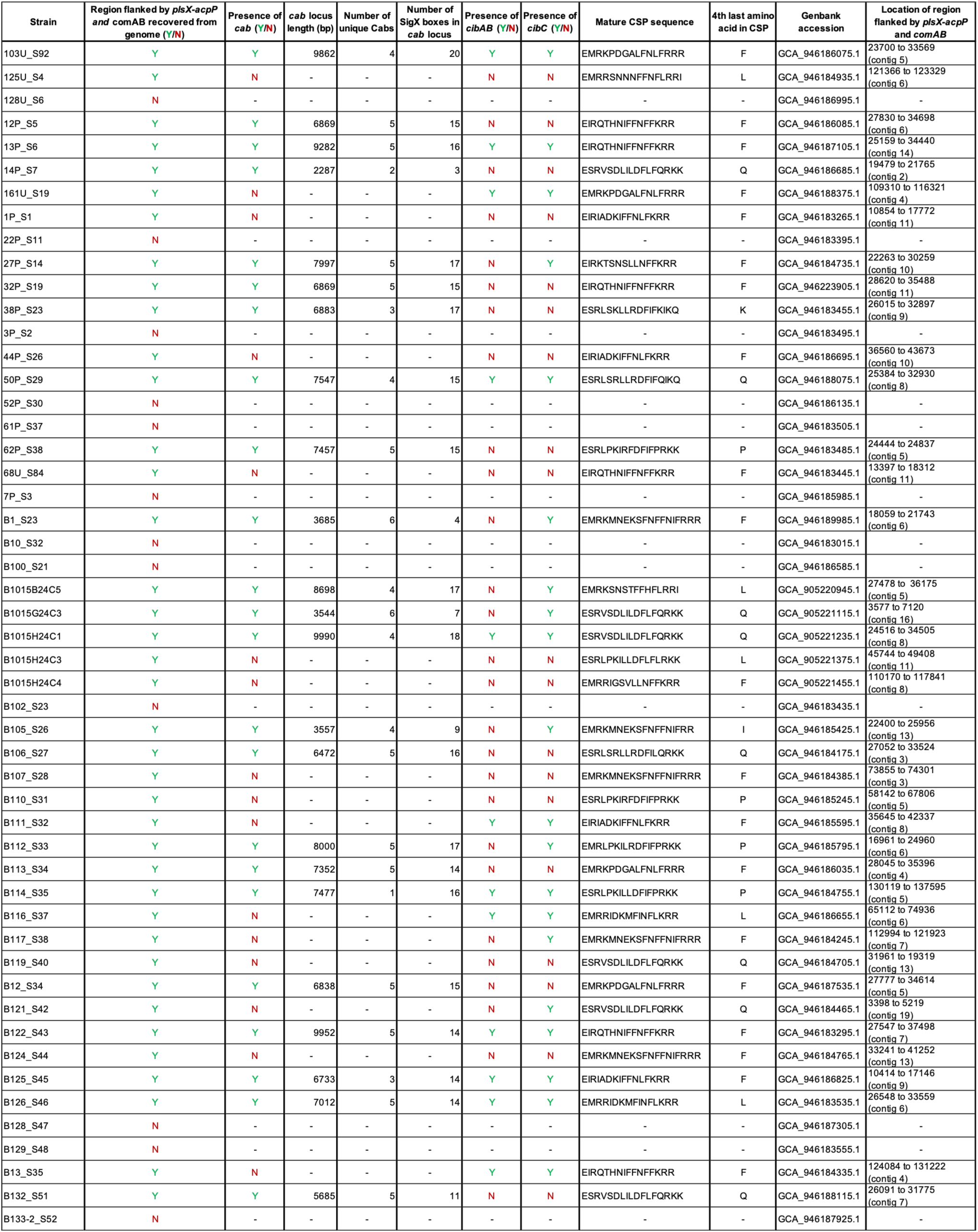
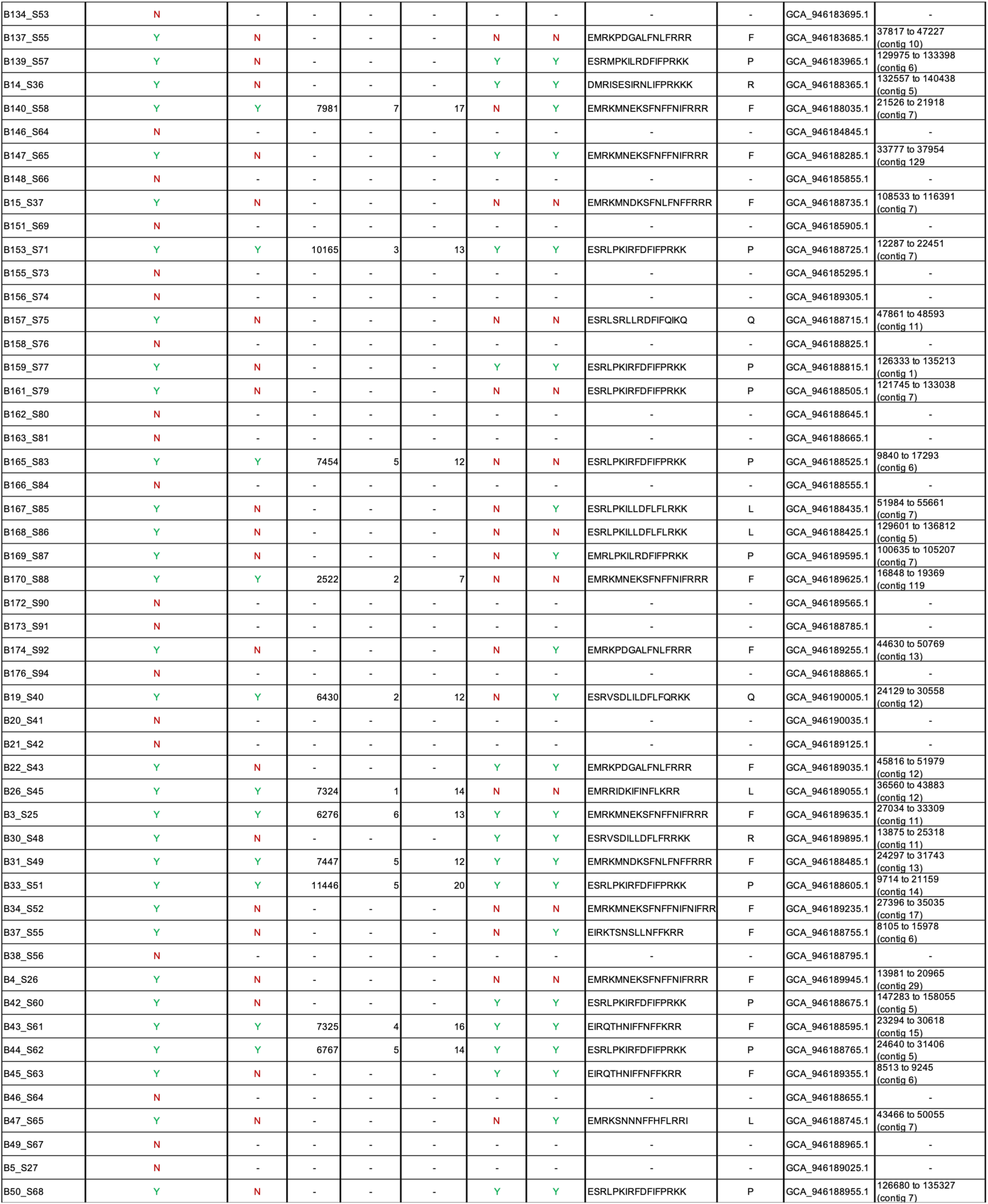
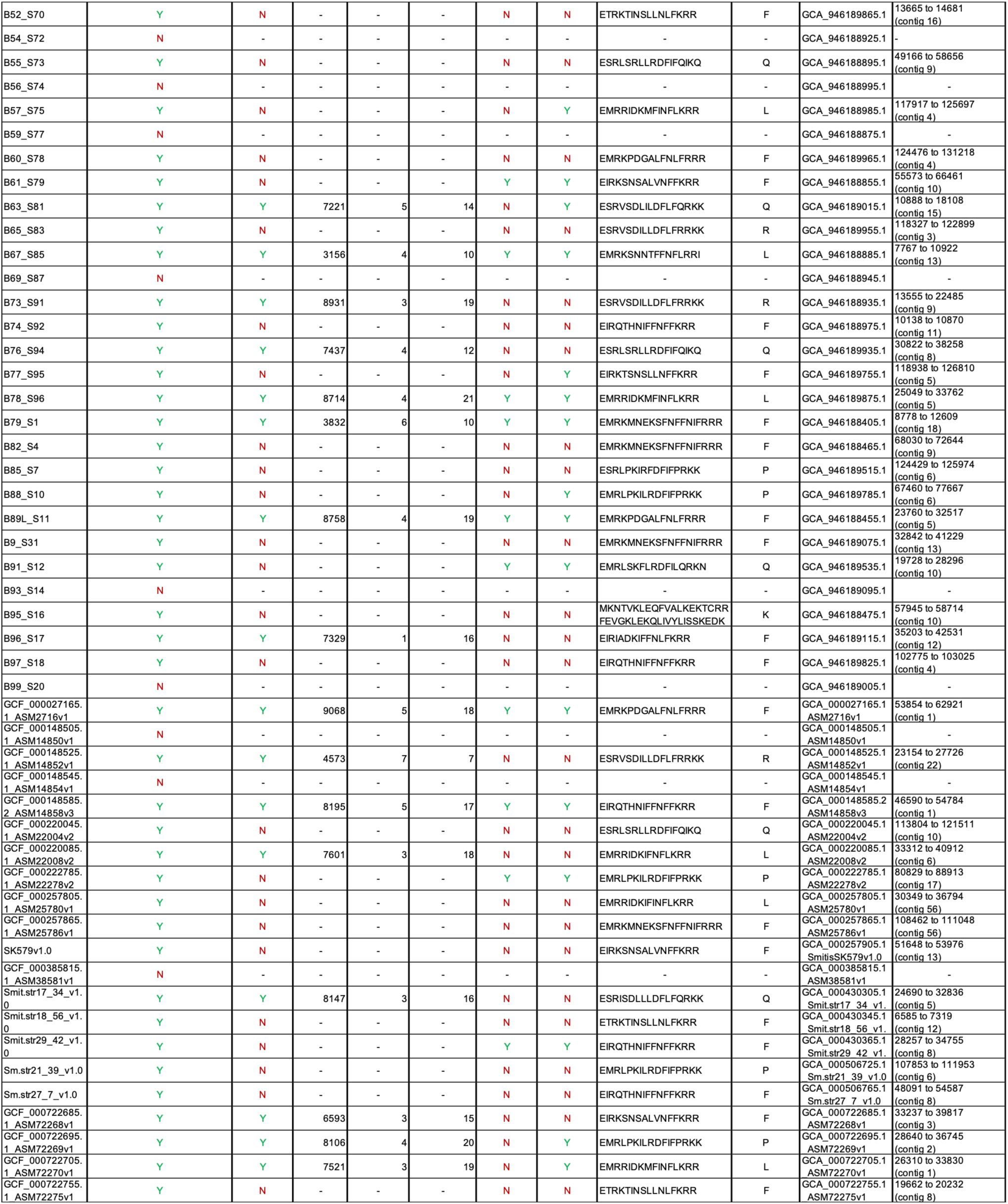
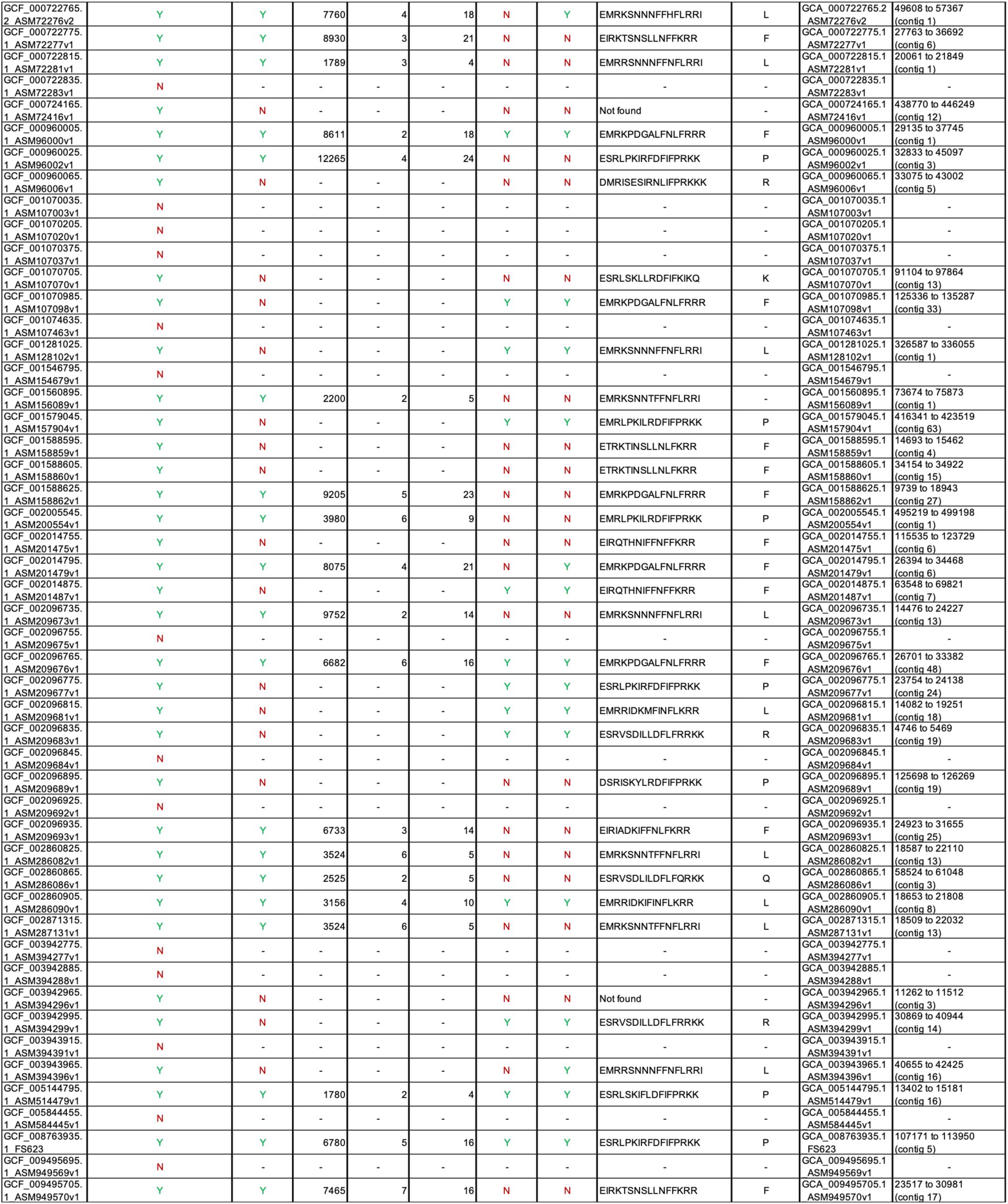
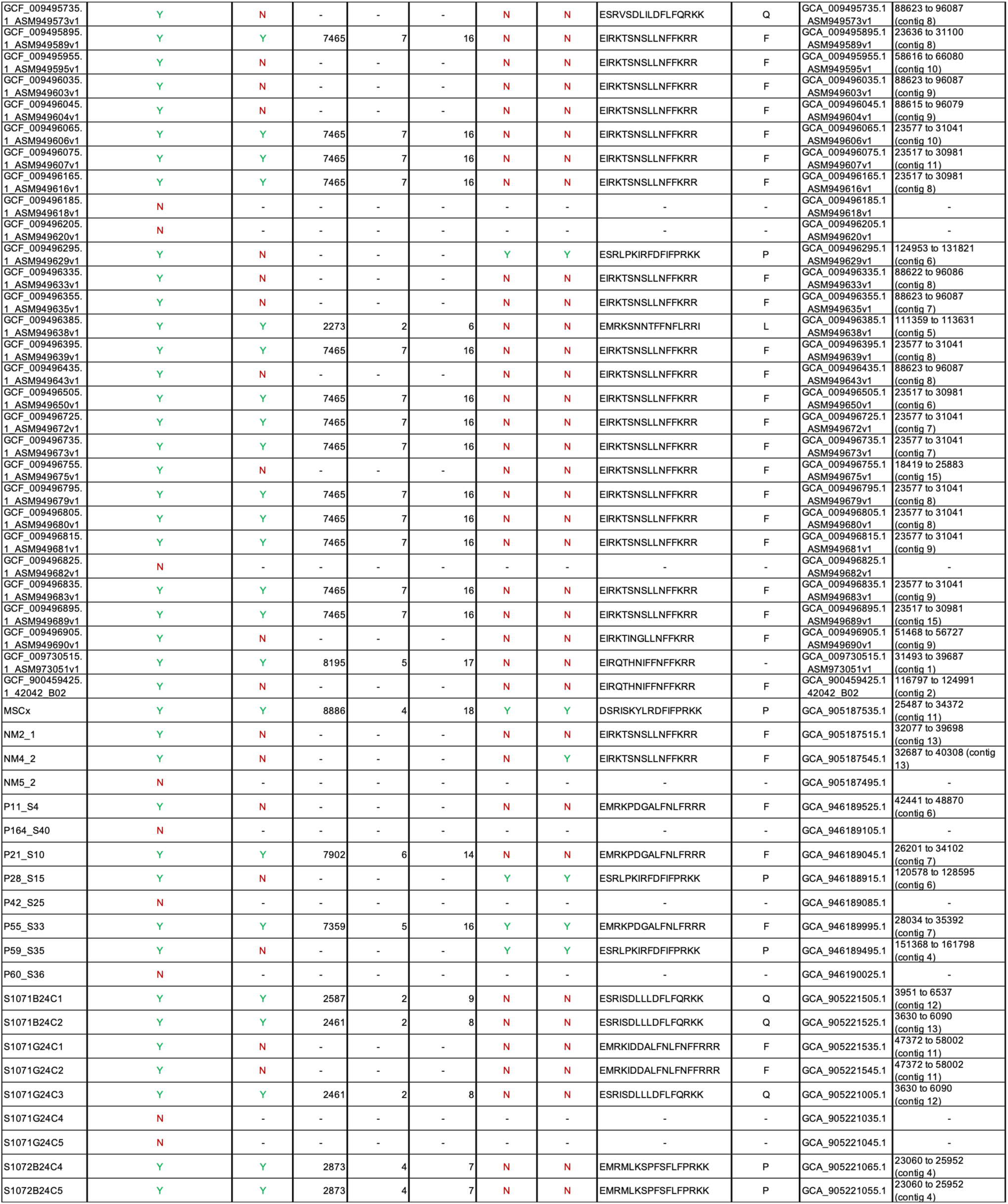
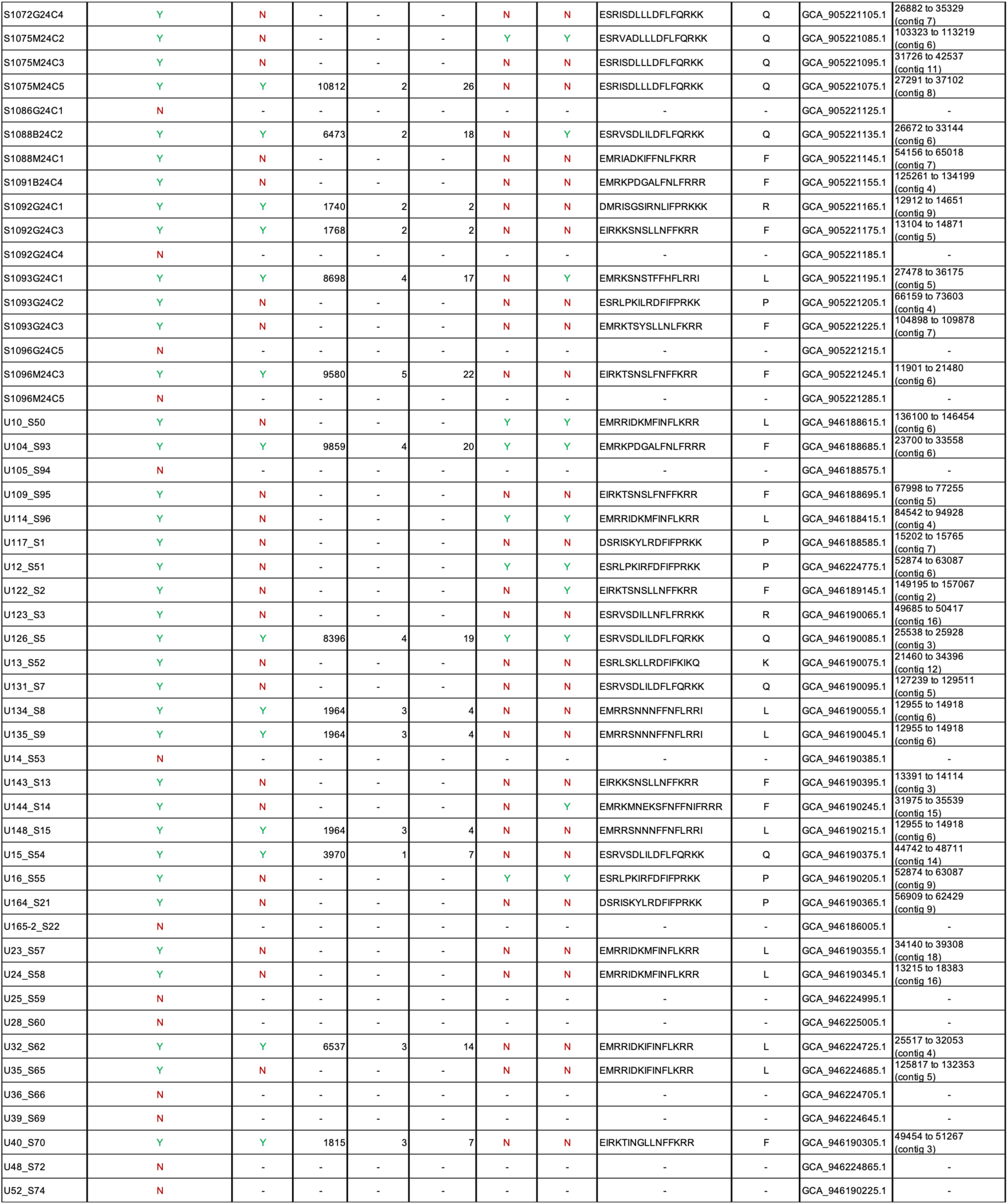
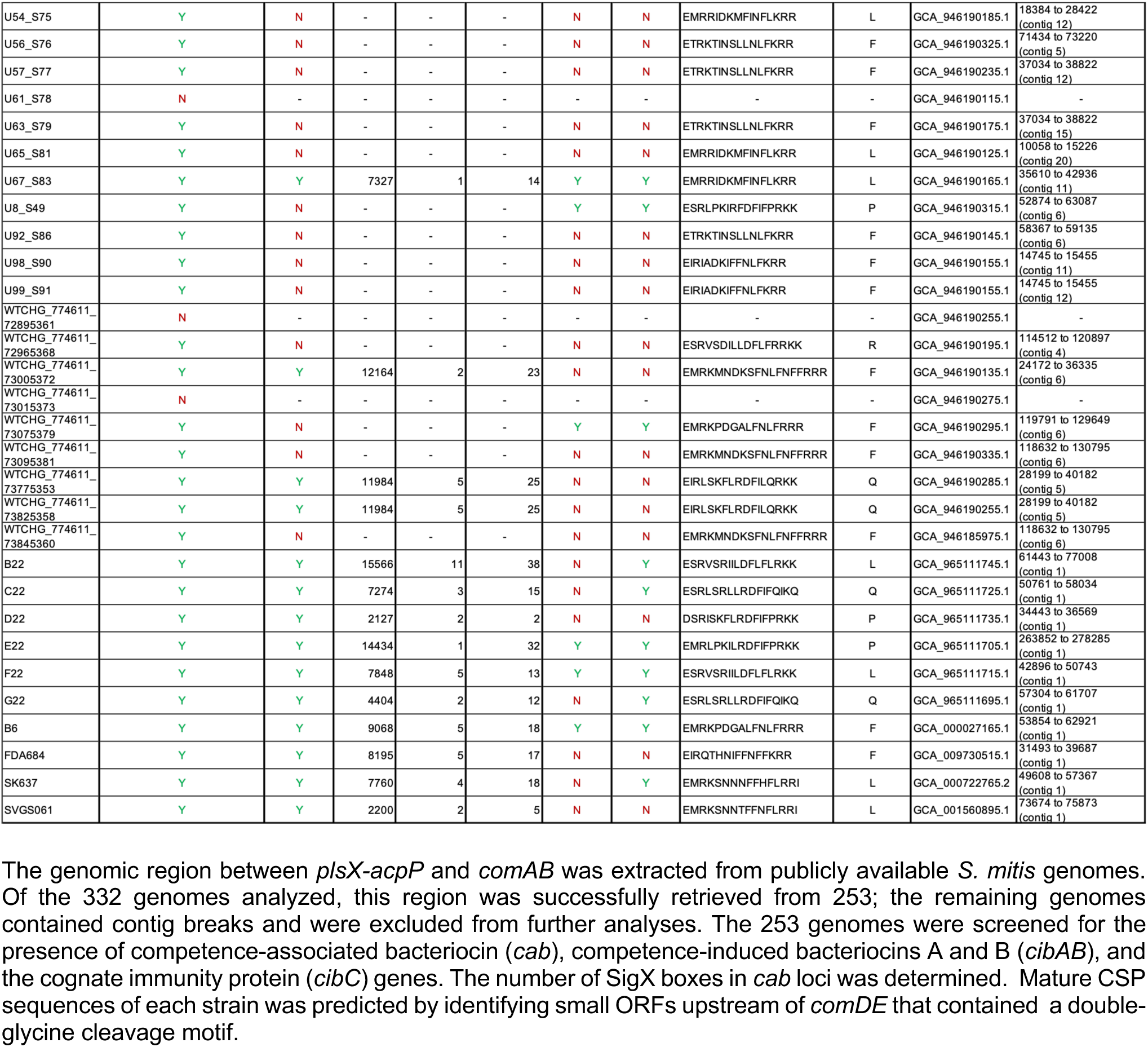
*Streptococcus mitis* genomes used in the study.

**Table S2.**
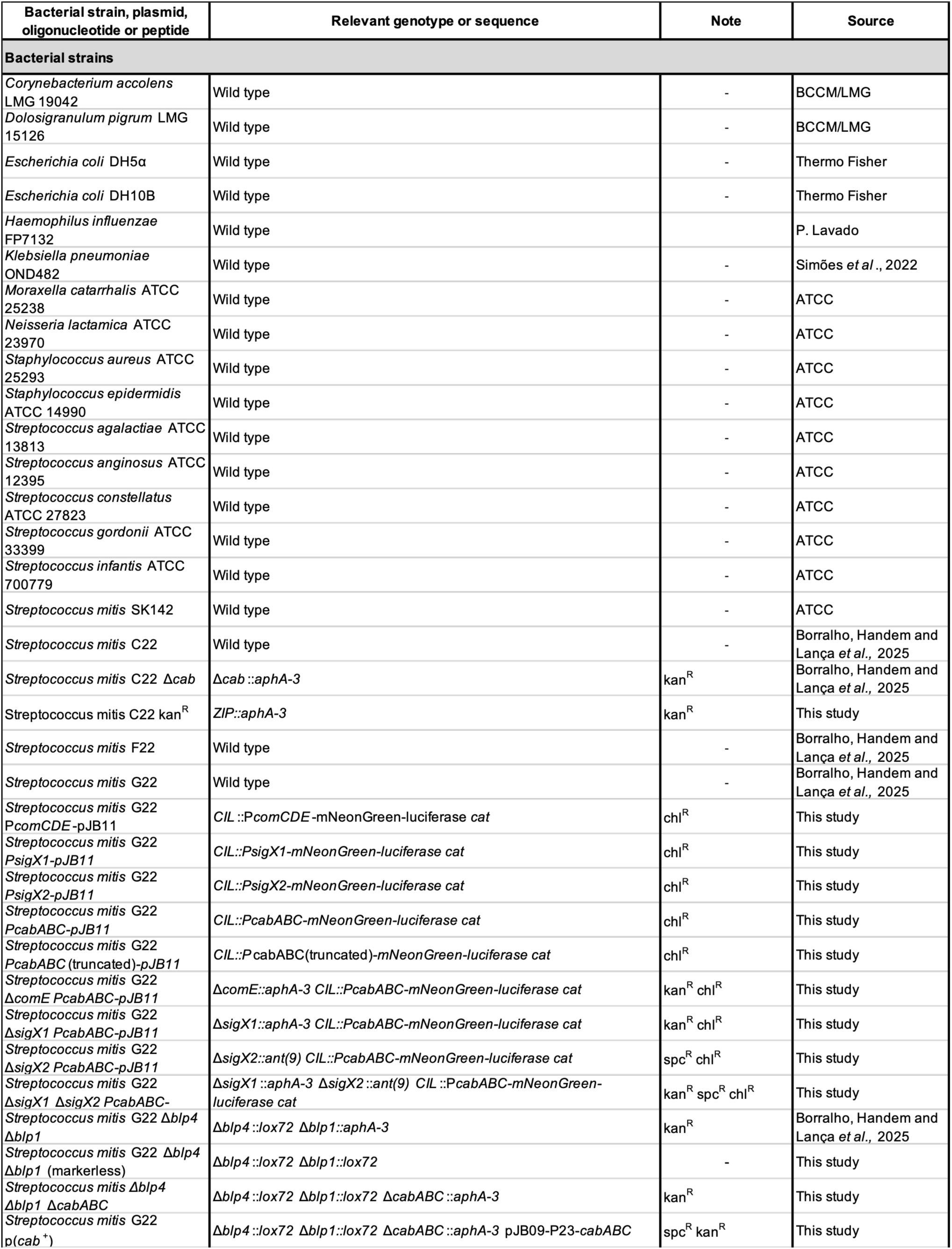
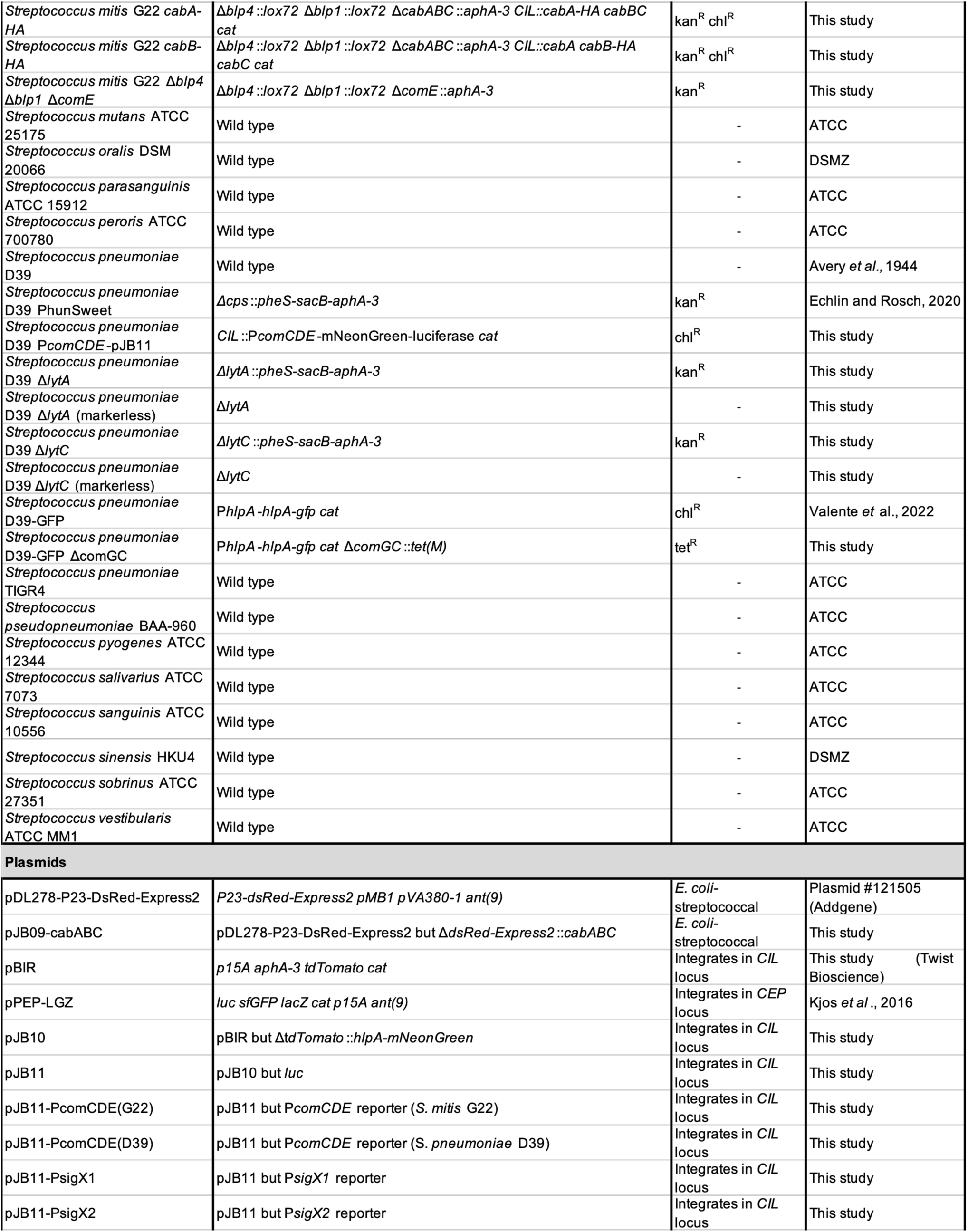
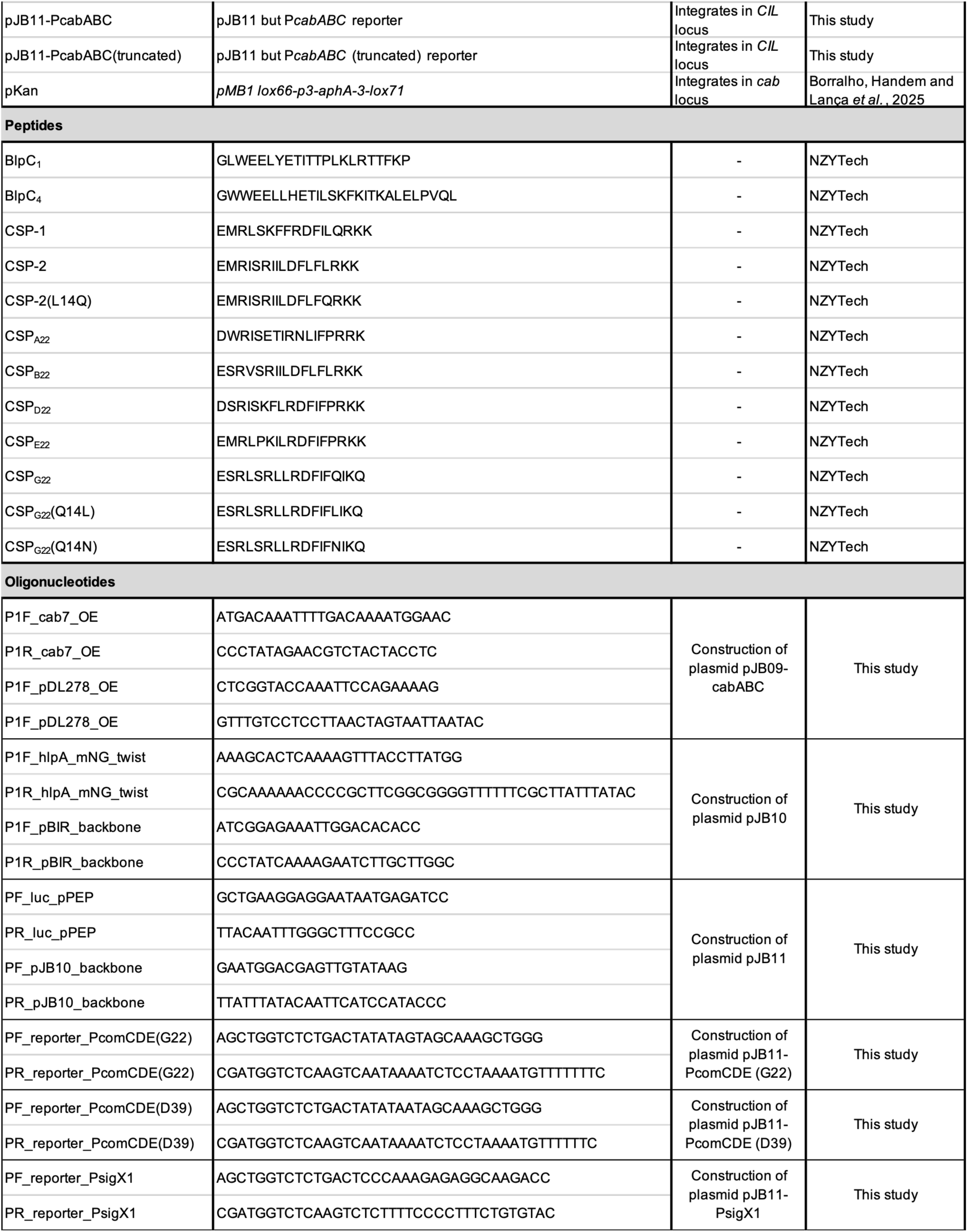
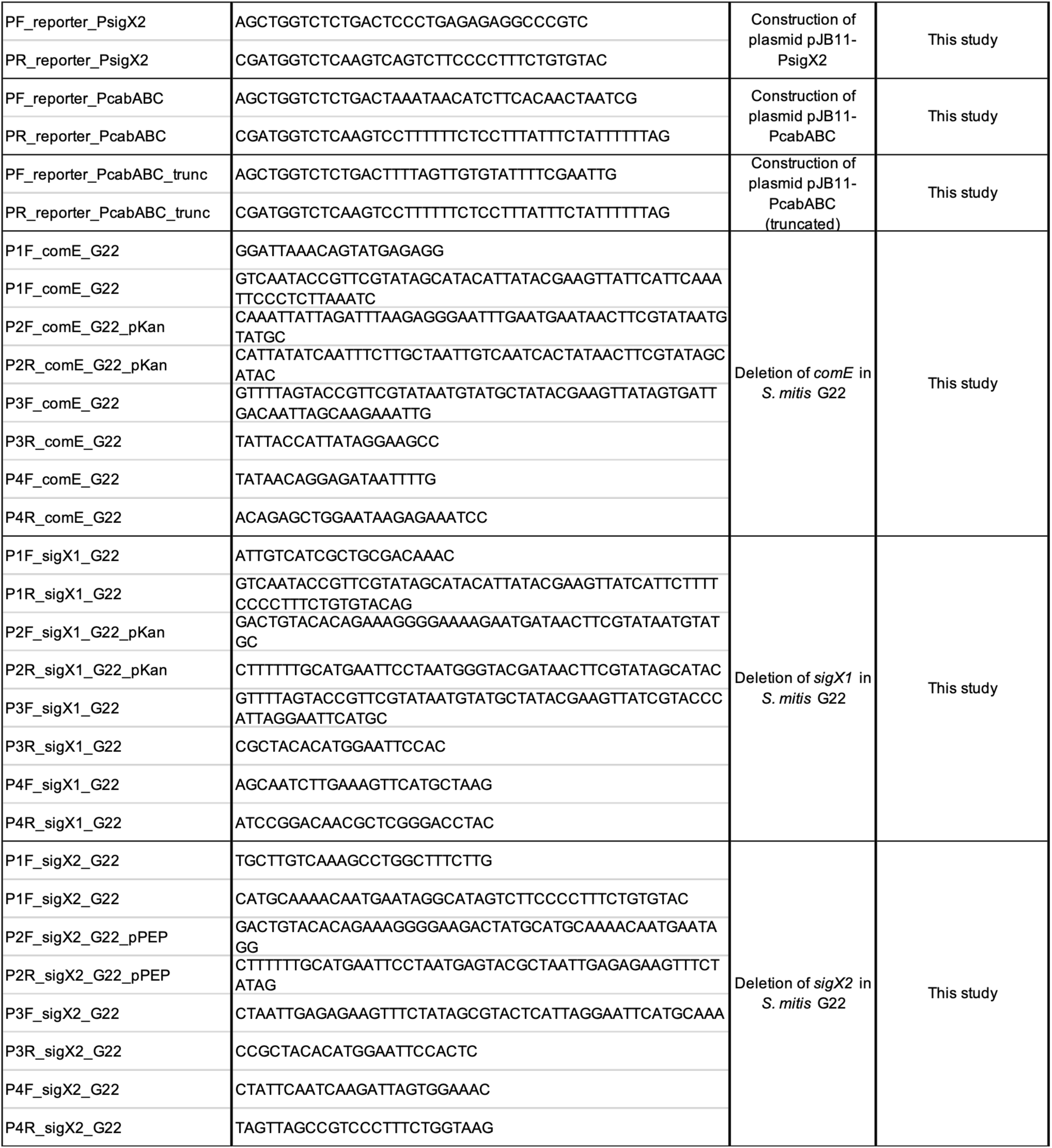
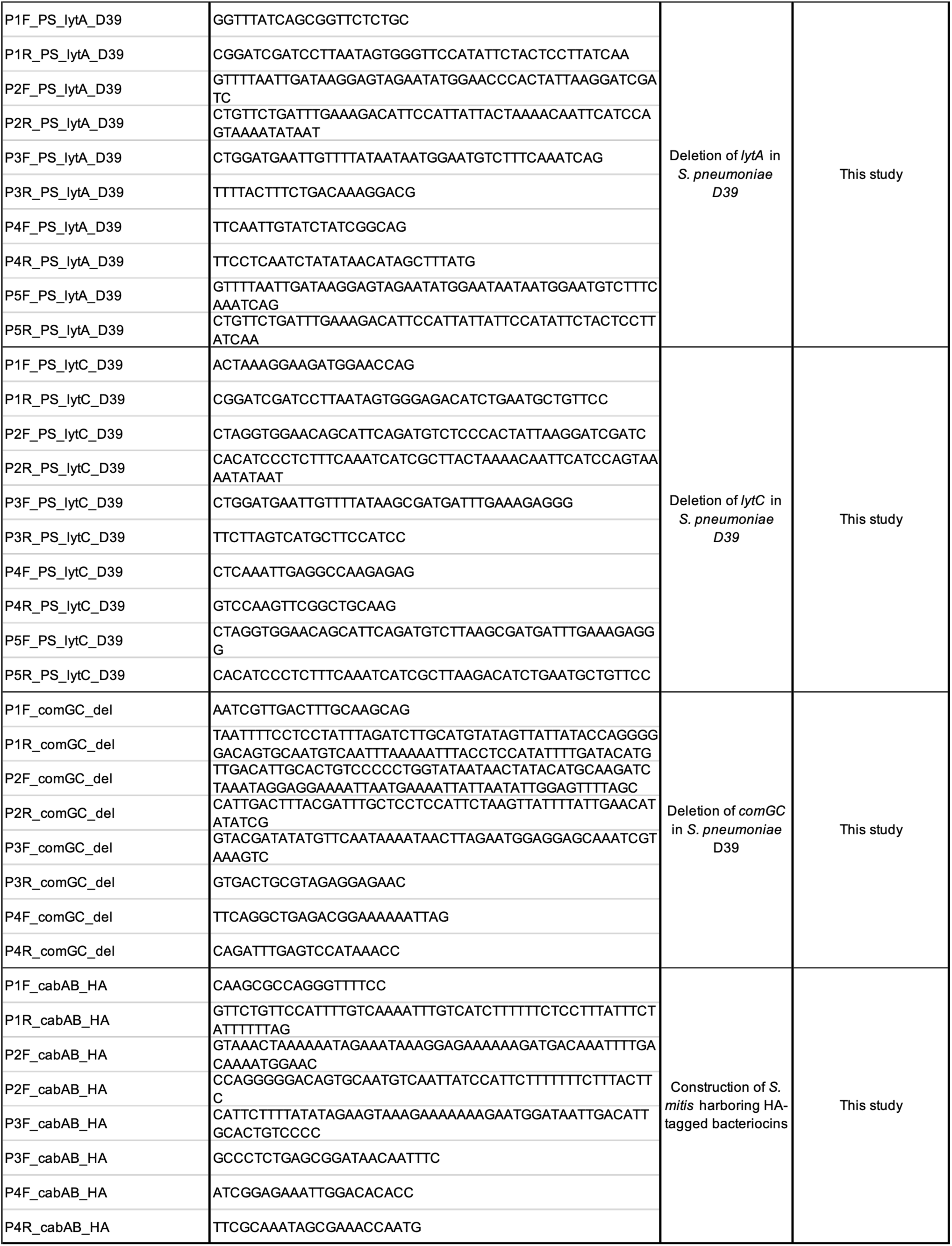
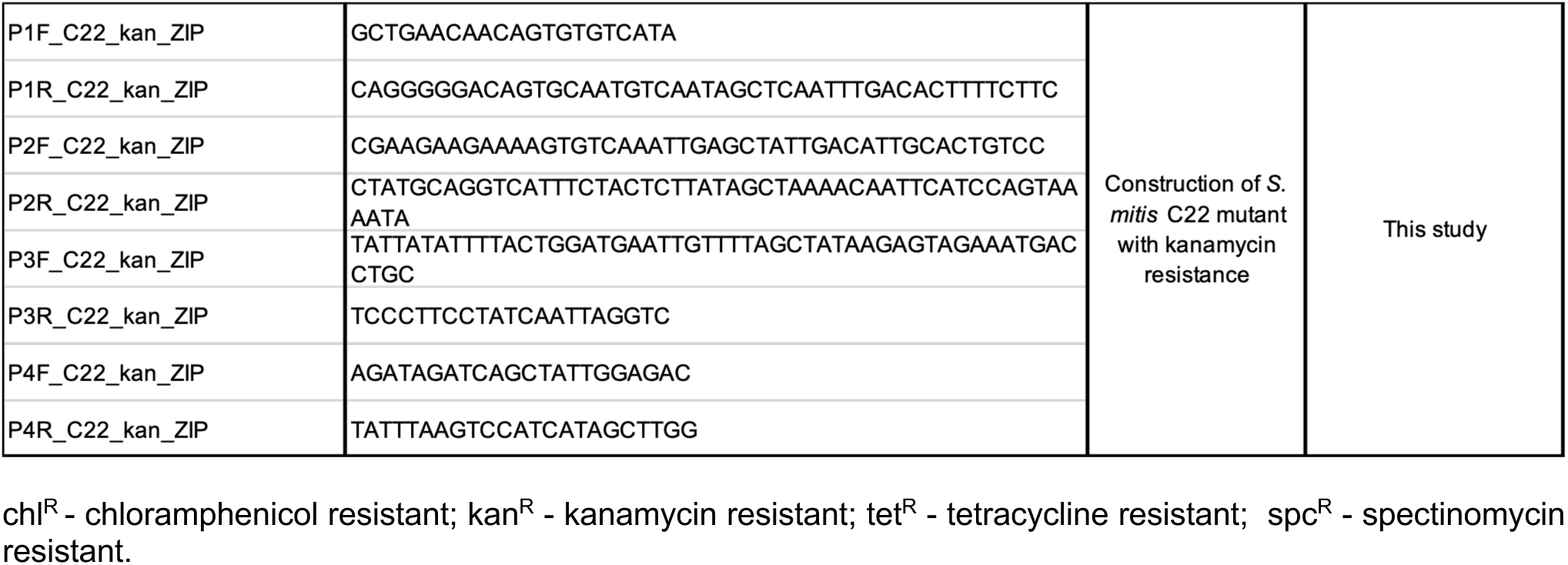
Bacterial strains, plasmids, peptides and oligonucleotides used in the study.

